# Deciphering Membrane Pore Formation Mechanisms of *Plasmodium falciparum* Perforin-Like Protein 1 (PfPLP1)

**DOI:** 10.1101/2025.10.14.682298

**Authors:** Sanket B. Patil, Subrata Dasgupta, Prasenjit Bhaumik

## Abstract

Membrane Attack Complex/Perforin (MACPF) domain proteins are β-pore forming toxins (β-PFTs) involved in the pathogenesis of various organisms. Among them, Perforin-like proteins (PLPs), produced by *Plasmodium* species, play essential roles in parasite invasion and egress. Due to increasing drug resistance in *Plasmodium*, PLPs represent promising but underexplored therapeutic targets, largely due to the lack of structural and mechanistic data. This study investigates the binding and pore formation mechanism of the *Plasmodium falciparum* PLP1 (PfPLP1), which is expressed during the human life cycle of the parasite. We modeled PfPLP1 structure and performed both all-atom and coarse-grained molecular dynamics simulations in soluble and membrane-associated states. PfPLP1 comprises two domains, a canonical MACPF domain and a β-pleated sheet domain-apicomplexan perforin β-domain (APCβ). Initial membrane binding is mediated by cationic residues at the base of the APCβ domain, which interact with the polar headgroups of the lipids from the host cell membrane. We analyzed the membrane-inserted tetrameric form where water molecules were observed to penetrate between the tetramer and the lipid bilayer, initiating pore opening. During this process, lipids reorganize into a toroidal edge to shield their hydrophobic tails, while water mixes with lipid headgroups in a disordered, heterogeneous fashion. Larger oligomeric assemblies show lateral displacement of lipids and a clear tendency to form pore-like structures. This study provides molecular insights into PfPLP1’s membrane binding and pore-forming behavior in both monomeric and oligomeric forms. The outcome of this study would be applicable in understanding pore formation mechanism in other PLPs and similar toxins.

**Graphical Abstract:** 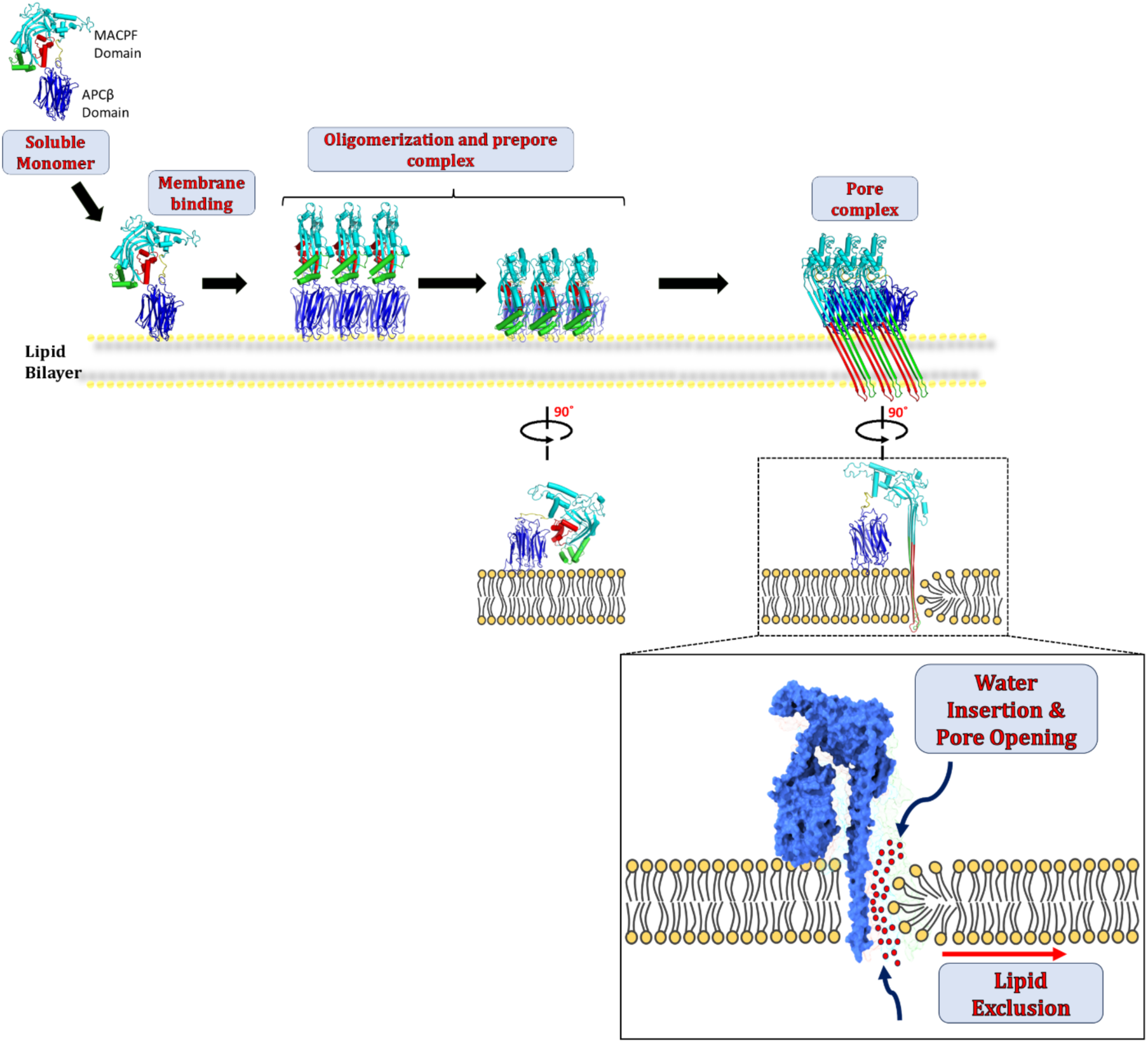

**Highlights:**

- Structure of *Plasmodium falciparum* PLP 1 (PfPLP1) as soluble form and membrane-inserted form.
- PfPLP1 initiates pore formation by binding to the host cell membrane with the help of the APC-β domain through cationic residues.
- Smaller oligomers of membrane-inserted PfPLP1 form an arc-like structure by reorganizing membrane lipids into a toroidal edge.
- Decamer and higher oligomers of membrane-inserted PfPLP1 form a pore-like structure by lateral displacement of the membrane lipids.

## Introduction

The plasma membrane of host organisms plays the first line of defense against any pathogenic infection. The pathogenic bacteria have developed toxins that target the plasma membrane to disrupt the integrity of the membrane through the formation of pores or channels (Gilbert, 2002). Pore-forming toxins (PFTs) are prevalent in bacteria, contributing to the virulence of pathogenic bacteria. The PFTs are also present in many other organisms, where PFTs are responsible for infection and attack, are involved in the invasion and egress of bacteria or parasites, and the defense system of the organism (Rosado *et al*., 2008; Reboul *et al*., 2016).

PFTs are classified into α-PFTs and β-PFTs based on the secondary structure that is responsible for the membrane insertion (Gouaux, 1997). Membrane attack complex/perforin (MACPF) and cholesterol-dependent cytolysins (CDC) proteins, which belong to the β-PFTs form one of the largest classes of PFTs. These MACPF/CDC family proteins possess a structurally conserved MACPF domain (Anderluh *et al*., 2014). Extensive structural and functional studies have been conducted on the monomeric forms of these proteins to understand their role in the pore formation mechanism. Recently, a few studies determined the structures of MACPF/CDC proteins in pre-pore and pore forms using single particle cryo-electron microscopy (cryo-EM) (Leung *et al*., 2014; Van Pee *et al*., 2017; Ni *et al*., 2020). Atomic force microscopy (AFM) and cryo-EM studies on CDC and MACPF proteins show the presence of membrane-inserted arcs (Sonnen *et al*., 2014; Leung *et al*., 2017). These results have shed lights on the mechanism of pore formation. However, the structural changes taking place during pore formation, which involve the conversion of helices present in the MACPF domain into the transmembrane β-hairpins (TMHs), the pore expansion, and the role of the membrane lipids, have still remained difficult to observe experimentally. Molecular dynamics simulations have been shown to help in understanding the dynamics of pore formation in both α-PFTs and β-PFTs ( Stoddart *et al*., 2014; Desikan *et al*., 2017a; Desikan *et al*., 2017b; Vögele *et al*., 2019; Desikan *et al*., 2020).

Apicomplexa parasites represent a diverse group of intracellular pathogens responsible for a myriad of diseases, ranging from malaria, caused by *Plasmodium* species like *Plasmodium falciparum, Plasmodium vivax*, to toxoplasmosis, attributed to *Toxoplasma gondii* (Kafsack and Carruthers, 2010). These parasites show the presence of proteins with the MACPF domain called Perforin-Like Proteins (PLPs) (Tavares *et al*., 2014). Five *Plasmodium* species are responsible for causing a life-threatening human disease called Malaria. In *Plasmodium*, five genes encoding PLPs have been identified, which are secreted during various stages of the parasite’s life cycle. These PLPs have a putative secretory signal sequence, suggesting that they are secreted proteins. These PLPs have an N-terminal segment, with unknown function, followed by a MACPF domain and a C-terminal domain called the apicomplexan perforin β (APCβ) domain (Kaiser *et al*., 2004). Various studies have shown that the PLPs are involved in the invasion and egress of the *Plasmodium* parasites (Ecker *et al*., 2007; Deligianni *et al*., 2013; Garg *et al*., 2013; Wirth *et al*., 2014). Recently, the crystallographic and molecular dynamics (MD) simulations studies were performed on only the APC-β domain structure of *P. vivax* PLP1 (PvPLP1) and PLP2 (PvPLP2) to understand the binding to lipid membrane (Williams *et al*., 2022). The APCβ domain of PvPLP2 has a membrane-interacting loop similar to the *T. gondii* APC-β domain (Guerra *et al*., 2018), having a conserved tryptophan residue responsible for the anchoring to the membrane. Although the importance of the PLPs is established for the invasion and egress of the *Plasmodium* parasite, there are no structural studies on the full-length PLPs. The interaction of the full-length protein with the membrane and the pore formation dynamics of the PLPs are yet to be understood.

In the present study, we have used computational approaches to investigate the pore formation mechanism of the *Plasmodium falciparum* PLP1 (PfPLP1). The All-atom and coarse-grained (CG) MD simulations were used to study interactions of PfPLP1 in monomeric form with the lipid membrane to analyse the binding of the protein. The monomeric PfPLP1 interacts with the lipid membrane with the Lysine and Arginine-rich loops present at the base of the protein. Simulations of the oligomeric arcs of PfPLP1 in the membrane-inserted form revealed that water molecules penetrate between the inner edge of the β-sheets formed by the oligomers and the lipid bilayer. This water insertion leads to pore expansion, while the lipid molecules at the pore edge rearrange to form a toroidal structure. Coarse-grained molecular dynamics simulations of larger membrane-inserted oligomers were also performed. The lipids are excluded laterally from the lumen of the oligomeric arcs, and the incomplete arcs form complete pore-like structures. The lipids present inside the complete oligomeric ring form a lipid plug. To the best of our knowledge, this is the first study demonstrating the interactions of PfPLP1 with the membrane and delineating the pore formation mechanism using oligomeric membrane-inserted PfPLP1.

## Materials and Methods

### Homology modeling of PfPLP1

The PfPLP1 structure was initially modeled using the AlphaFold2 computational program (Jumper *et al*., 2021). The domains were modeled well for the soluble form of PfPLP1; however, the arrangement of the domains was unlikely. The AlphaFold2 and AlphaFold3 (Abramson *et al*., 2024) were unable to predict the structure of the PfPLP1 in membrane-inserted form. Thus, we used template-based homology modeling for the PfPLP1 structure in soluble and membrane-inserted form. The uncharacterized N-terminal segment (0-238 aa) was excluded from the structure modeling. The SWISS-MODEL (Kiefer *et al*., 2009) server was unable to generate the full-length model of PfPLP1 on the basis of a single template. Therefore, two different templates were used to build the two domains of the model. Two different templates were used to model the MACPF domain (239-564 aa) in two forms: soluble and membrane-inserted form. The cryo-EM structures of the murine perforin-2, which showed a 25.77% identity with PfPLP1, were used as a template for modeling the MACPF domain (Ni *et al*., 2020). The murine perforin2 pre-pore structure (PDB ID: 6SB3) was used for modeling the MACPF domain in soluble form, and the murine perforin2 in pore structure (PDB ID: 6SB5) was used for the membrane-inserted form. The X-ray crystallographic structure of the perforin-like protein 1 from *Toxoplasma gondii* (TgPLP1) (PDB ID: 6D7A), which showed 28.91% identity to PfPLP1, was used as the template for the APCβ domain (580 – 840 aa) (Guerra *et al*., 2018). The two domains were connected by Modeller 10.2 with the linker loop (565 – 579 aa) to get the final soluble PfPLP1 and membrane-inserted PfPLP1 structure (Eswar *et al*., 2008).

### Simulation system preparation

The soluble PfPLP1 monomer was solvated with a cubic water box of TIP3P water molecules with 10 Å from any edge of the protein surface, and the entire system was neutralized with an appropriate number of Na^+^ and Cl^-^ ions. The system was used for an all-atom MD simulation of PfPLP1 in soluble form to mimic its physiological state during secretion in monomeric form. The soluble PfPLP1 and membrane-inserted PfPLP1 were docked separately on a lipid bilayer composed of POPC (45%), POPE (45%), and POPS (10%) using the CHARMM-GUI server (membrane builder module) for all-atom MD simulations (Huang and MacKerell Jr, 2013; Wirth *et al*., 2014; Williams *et al*., 2022). Both the systems were further solvated with TIP3P water molecules till 60 Å on both sides of the membrane and neutralized with an appropriate number of Na^+^ and Cl^-^ ions. The systems were designed to mimic the physiological binding of soluble PfPLP1 to the lipid membrane, as well as its subsequent insertion into the bilayer. A similar setup was used for the all-atom simulation of the membrane-inserted PfPLP1 tetramer. The tetrameric protein assembly was generated using the cryo-EM structure of murine perforin 2 pore structure (PDB ID: 6SB5) with COOT, which represents the incomplete arcs (Emsley *et al*., 2010). As the oligomeric system was becoming complex with the number of monomer subunits in the system, coarse-grained MD simulations were performed for larger oligomeric arcs. Membrane-inserted form of PfPLP1 in hexamer, octamer, decamer, dodecamer, tetradecamer, and hexadecamer (complete ring) assemblies were generated similarly to that of the tetramer assembly. The Martini maker (bilayer builder) module of the CHRAMM-GUI (Jo *et al*., 2008) server was used for generating the systems, and the martini22P (Qi *et al*., 2015) model was used for all the systems. The composition of the lipid bilayer membrane was kept the same as mentioned above, and the dimensions were adjusted to accommodate the complete assembly of oligomers.

### All-atom molecular dynamics simulations

The molecular dynamics (MD) simulations of the protein structures was conducted using GROMACS v2018.8 with the CHARMM36 force field (Huang and MacKerell Jr, 2013). Energy minimization was carried out using the steepest descent method, applying 50,000 steps for all systems with a tolerance of 1000 kJ mol^−1^ nm^−1^. Equilibration was performed for 100 for each system under constant particles, volume, and temperature (NVT, using a modified Berendsen thermostat (Berendsen *et al*., 1984) with velocity rescaling at 310K and a 0.1 ps time step) and under constant particles, pressure, and temperature (NPT, utilizing Parrinello−Rahman pressure coupling (Parrinello and Rahman, 1981) at 1 bar with a compressibility of 4.5 × 10^−5^ bar^−1^ and a 2 ps time constant). The Nosé−Hoover thermostat (Nosé, 1984; Hoover, 1985) was used during NPT equilibration and the final production run to ensure proper kinetic ensembles. Bond lengths were constrained with the Linear Constraint Solver (LINCS) algorithm. For long-range interactions, the particle-mesh Ewald (PME) method was employed with a 1.2 nm cutoff and a Fourier spacing of 0.16 nm. After equilibration, the systems underwent a 100 ns production run, with trajectories saved every 10 ps for data analysis.

### Coarse-grained MD-simulation

The coarse-grained (CG) models have been proven to be very useful and feasible with a simulation time scale and produce data comparable to the all-atom models (Kmiecik *et al*., 2016). CG simulation is advantageous with large macromolecular complexes, particularly the lipid-protein complexes. The coarse-grained models PfPLP1, consisting of hexamer, octamer, decamer, dodecamer, tetradecamer, and hexadecamer on a lipid bilayer in membrane-inserted form, were built with the help of the CHARMM-GUI Martini Maker (Qi *et al*., 2015), which allows automated construction of coarse-grained models based on the Martini 22P force field. The atomic structure of the target membrane protein was first converted to a coarse-grained representation using CHARMM-GUI (Jo *et al*., 2008). A lipid bilayer was constructed around the protein using the desired lipid composition (POPC, POPE, and POPS), and the system was solvated with Martini water beads and neutralized with appropriate counter ions. Standard Martini 3 parameters were used for protein, lipids, water, and ions. The system was energy-minimized using the steepest descent algorithm to remove any unfavourable contacts. Following minimization, a multi-step equilibration protocol was carried out using position restraints on the protein and lipids to allow the solvent and membrane to relax. The equilibration was followed by production runs under NPT conditions. Temperature was maintained at 310 K using the velocity-rescaling thermostat (Bussi *et al*., 2007) with a coupling constant of 1.0 ps, and pressure was maintained semi-isotropically at 1 bar using the Parrinello-Rahman barostat (Parrinello and Rahman, 1981) with a coupling constant of 12.0 ps and compressibility of 3×10^-4^ bar^-1^. The time step for integration was set to 20 fs, and periodic boundary conditions were applied in all three dimensions. Non-bonded interactions were treated with a cutoff of 1.1 nm, and the neighbor list was updated every 20 steps. Each simulation was run for a total of 1 μs, and data were collected at appropriate intervals for subsequent analysis.

### Data analysis

The simulation trajectories were visualized using VMD (Humphrey *et al*., 1996), and PyMOL (Schrödinger L, 2015). Analyses included RMSD, RMSF, lipid–protein interactions, and membrane deformation, using standard GROMACS tools (Lindahl *et al*., 2001), and custom Python scripts.

## Results and Discussion

### Structural fold of PfPL1

The full-length sequence of PfPLP1 (UniProt ID: Q9U0J9) with 842 residues was retrieved from the UniProt database. The full-length protein has three distinct parts called the N-terminal segment, followed by the MACPF domain, and the C-terminal APCβ domain (Figure 1A). The N-terminal segment consists of 238 amino acids, which also contains the putative signal peptide. The function of the N-terminal segment is not well characterized (Kaiser *et al*., 2004). The model of the peptide containing residues 239 to 840, excluding the N-terminal segment, was generated in soluble form (Figure 1B) and in membrane-inserted form (Figure 1C).

**Figure 1.**
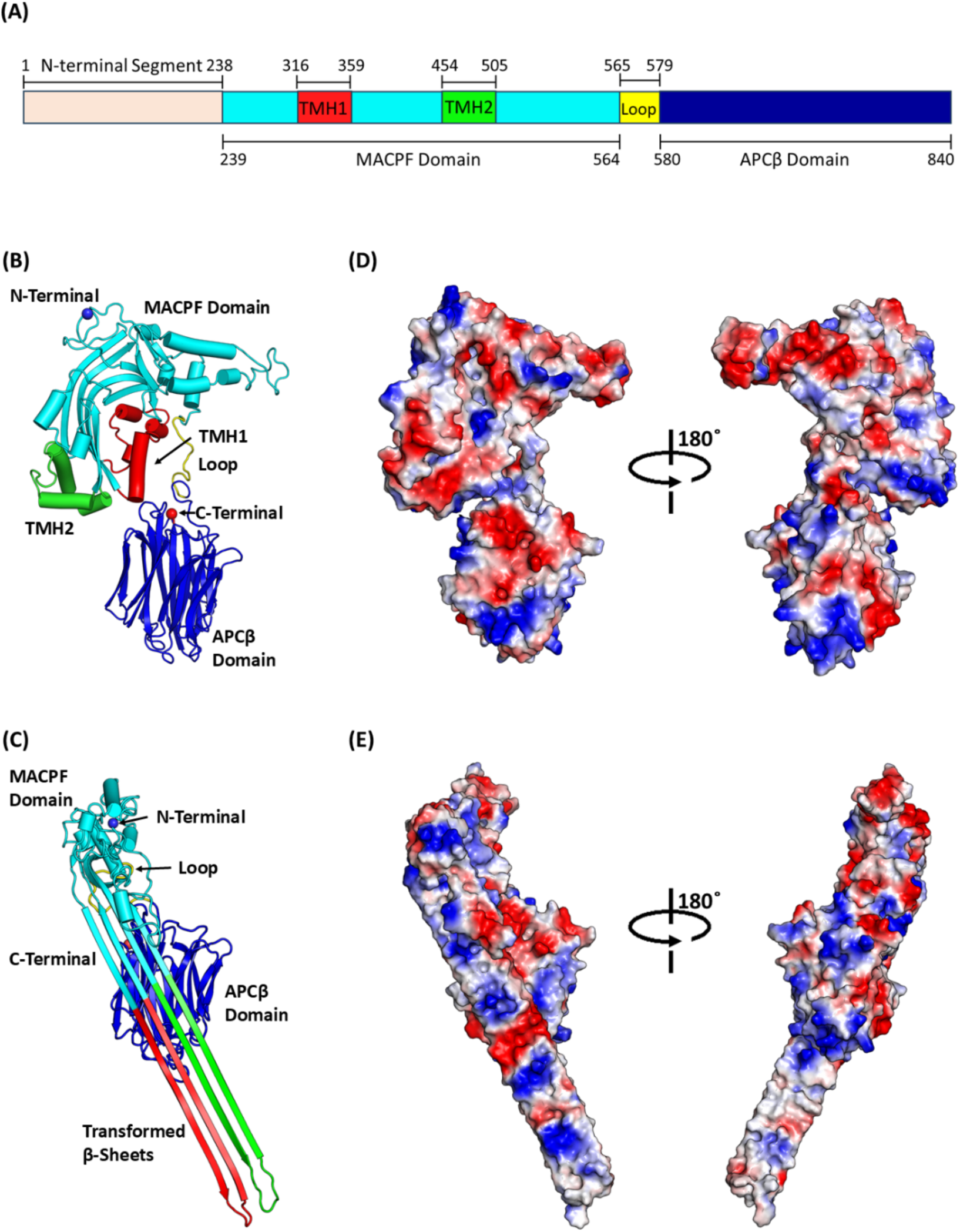
The domain architecture and overall structural fold of the modeled PfPLP1 monomer. (A) The bar diagram shows the domain architecture of the full-length PfPLP1 polypeptide. The light pink bar represents the N-terminal segment, the cyan bar represents the MACPF domain, and the blue bar represents the APCβ domain. The TMHs present in the MACPF domain are represented in red (TMH1) and green (TMH2). The loop connecting the two domains is highlighted in yellow. (B) The cartoon representation of modeled PfPLP1 in soluble form. (C) The cartoon representation of the modeled PfPLP1 membrane-inserted form. The colors for the domains are kept the same as the bar diagram for better understanding. (D) The electrostatic surface charge distribution (blue: positive, red: negative, and grey: neutral) of PfPLP1 in soluble form. (E) The electrostatic surface charge distribution (blue: positive, red: negative, and grey: neutral) of PfPLP1 in membrane-inserted form.

The stereochemical quality of the modeled structure was verified using the PROCHECK server (Laskowski *et al*., 1993). The Ramachandran plot analysis showed that 96.3% of residues are present in the allowed region of the map for soluble PfPLP1 (Figure S1A), and 97 % of residues were present in the allowed region of the map for the membrane-inserted form of PfPLP1 (Figure S1B). The MACF domain is composed of α-helices and β-sheets, whereas the APCβ domain is entirely composed of β-sheets joined by random coils. The MACPF domain contains 326 amino acids, ranging from 239 to 564, while the APCβ domain is made from 261 amino acids, ranging from 580 to 840. The two domains are linked by a 15-residue loop (565–579). The MACPF domain features central β-strands flanked by helical segments, which transition into transmembrane β-hairpins (TMHs) upon pore formation. TMH1 and TMH2 span residues 316–359 and 454–505, respectively (Figure 1A). During membrane insertion, these β-strands and TMHs rearrange to contribute to the formation of the transmembrane β-sheet structure, facilitating pore assembly.

The surface charge distribution of PfPLP1 shows the presence of amphipathic residues in the MACPF domain, especially in the TMHs region (Figure 1D). The TMHs have alternate polar and non-polar residues, which can be seen in the surface representation of the membrane-inserted form. In this conformation, one face of the β-sheets in the membrane-inserted form mostly contains charged residues (Figure 1E left side). In contrast, the other side is mostly composed of non-polar residues (Figure 1E right side), suggesting an amphipathic organization conducive to membrane interaction and insertion. These modeled structures were further used to understand the pore formation mechanism of PfPLP1.

### Evaluation of the stability of PfPLP1 in water and on lipid bilayer

The stability of the PfPLP1 monomer was evaluated by molecular dynamics (MD) simulations, both in aqueous solution and when bound to a lipid bilayer. The stability assessment was done based on the Root Mean Square Deviation (RMSD) of the backbone atoms and the radius of gyration (Rg) of PfPLP1. Three systems were analysed: soluble form in solvent (Figure 2A), soluble form on lipid bilayer (Figure 2B), and the membrane-inserted form on lipid bilayer (Figure 2C), for which the normalized RMSD and Rg values were plotted, and shown in Figures 2D, 2E, 2F, for three systems respectively. During the 100 ns simulation run, the soluble PfPLP1 in solvent took around 25ns to stabilize. Once the structure was stabilized, the normalized RMDS values ranged between 0.78 – 0.96 units. The soluble PfPLP1 on lipid bilayer and membrane-inserted PfPLP1 on lipid bilayer systems were stabilized comparatively faster, within 15 ns. The normalized RMSD values for the soluble PfPLP1 on lipid bilayer and membrane-inserted PfPLP1 on lipid bilayer range between 0.75-0.95 units and 0.62-0.9 units. The soluble PfPLP1 monomer in solvent exhibits a higher RMSD, indicating greater deviation from the initial structure and suggesting significant conformational changes. These changes are most pronounced in the TMH helices in the soluble state. The same region adopts a β-sheet conformation and is inserted into the lipid bilayer; it exhibits a minimal deviation, indicating enhanced stability of the region in the membrane environment. Additionally, the radius of gyration remains consistent across all three PfPLP1 monomer systems and is stable throughout the simulation, suggesting that the overall structure remains compact. All the time-dependent simulation curves are shown in the supplementary figure (Figure S2).

**Figure 2:**
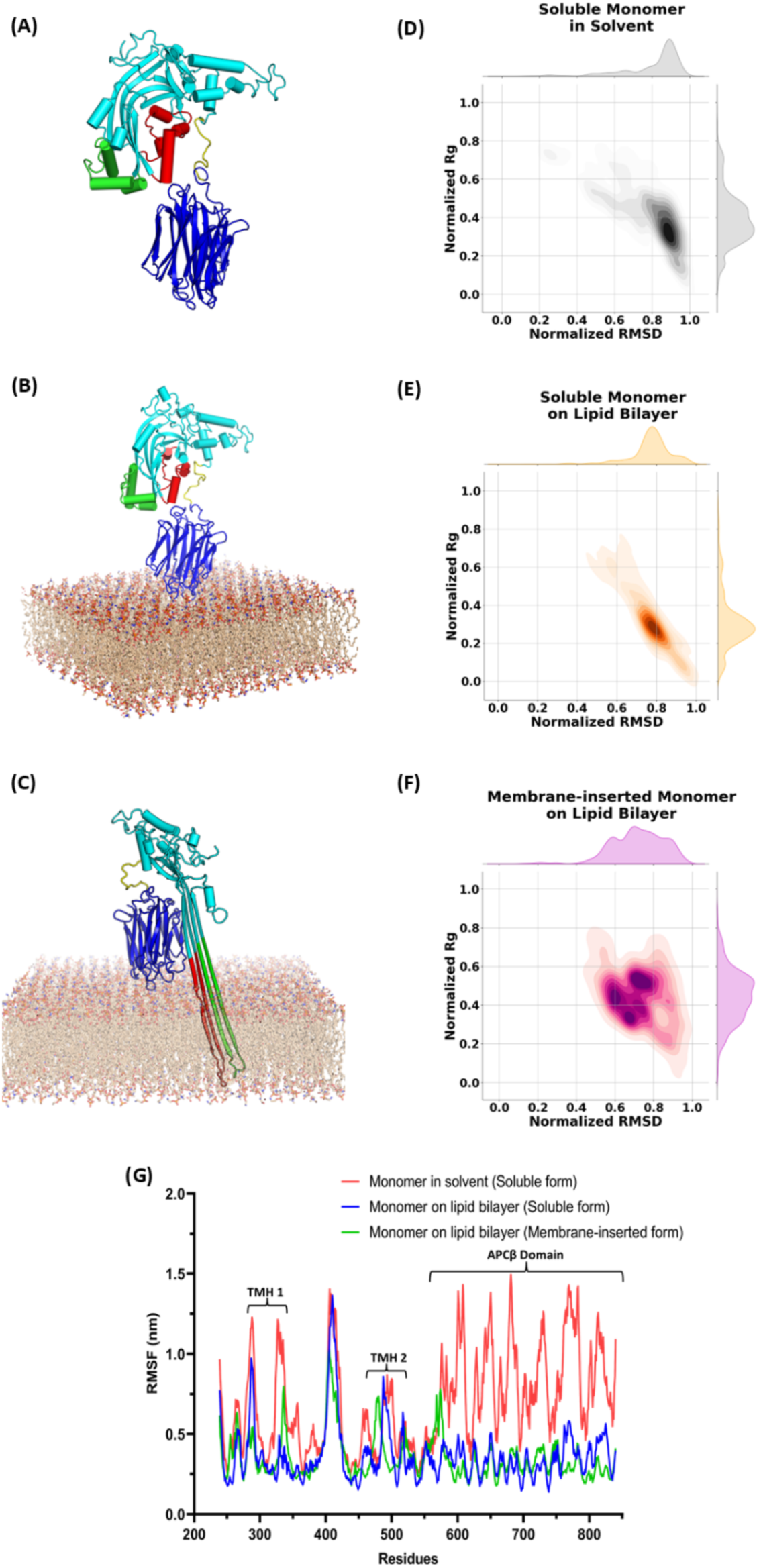
All-atom simulations of PfPLP1 monomer, the Root Mean Square Deviation (RMSD) vs Radius of Gyration (Rg) plots, and the Root Mean Square Fluctuation (RMSF) plot. Three systems were simulated: (A) Soluble PfPLP1 in solvent, (B) Soluble PfPLP1 on lipid bilayer, (C) Membrane-inserted PfPLP1. (D) Normalized RMSD vs Rg plot for soluble PfPLP1 in solvent. (E) Normalized RMSD vs Rg plot for soluble PfPLP1 on lipid bilayer. (F) Normalized RMSD vs Rg plot for membrane-inserted PfPLP1 on lipid bilayer. Different forms of PfPLP1 are presented as a cartoon (colors for the domains are the same as mentioned in Figure 1) and the lipid bilayer as a stick model. (G) The RMSF values for all three systems are shown in different colors: the soluble PfPLP1 in solvent (red), the soluble PfPLP1 on lipid bilayer (blue), and the membrane-inserted PfPLP1 on lipid bilayer (green).

The Root Mean Square Fluctuations (RMSF) indicate how much amino acid residues deviate from their average positions during simulations. RMSF plots for the three systems are presented in the Figure. 2G. The APCβ domain exhibits greater fluctuations in the solvent, with values ranging from 0.4 to 1.5 nm. In both the monomer on the lipid bilayer and the membrane-inserted forms of PfPLP1, these fluctuations are reduced to between 0.2 and 0.5 nm. Similarly, the TMH helices exhibit greater fluctuations in the solvent phase, while the membrane-inserted form displays the lowest fluctuations upon formation of β-sheets, suggesting enhanced structural stability upon membrane association.

The base of the APCβ domain, which interacts with the lipid bilayer, is populated with positively charged residues (Figure 1D, 1E). The PfPLP1 anchors to the lipid membrane with the 6 loops at the base of the APCβ domain. The loops (called here, L1-L6) are highlighted with different colors (Figure 3). These loops are characterized by the presence of positively charged residues (Lys and Arg) and some polar residues like Ser, which interact with the lipid bilayer.

**Figure 3:**
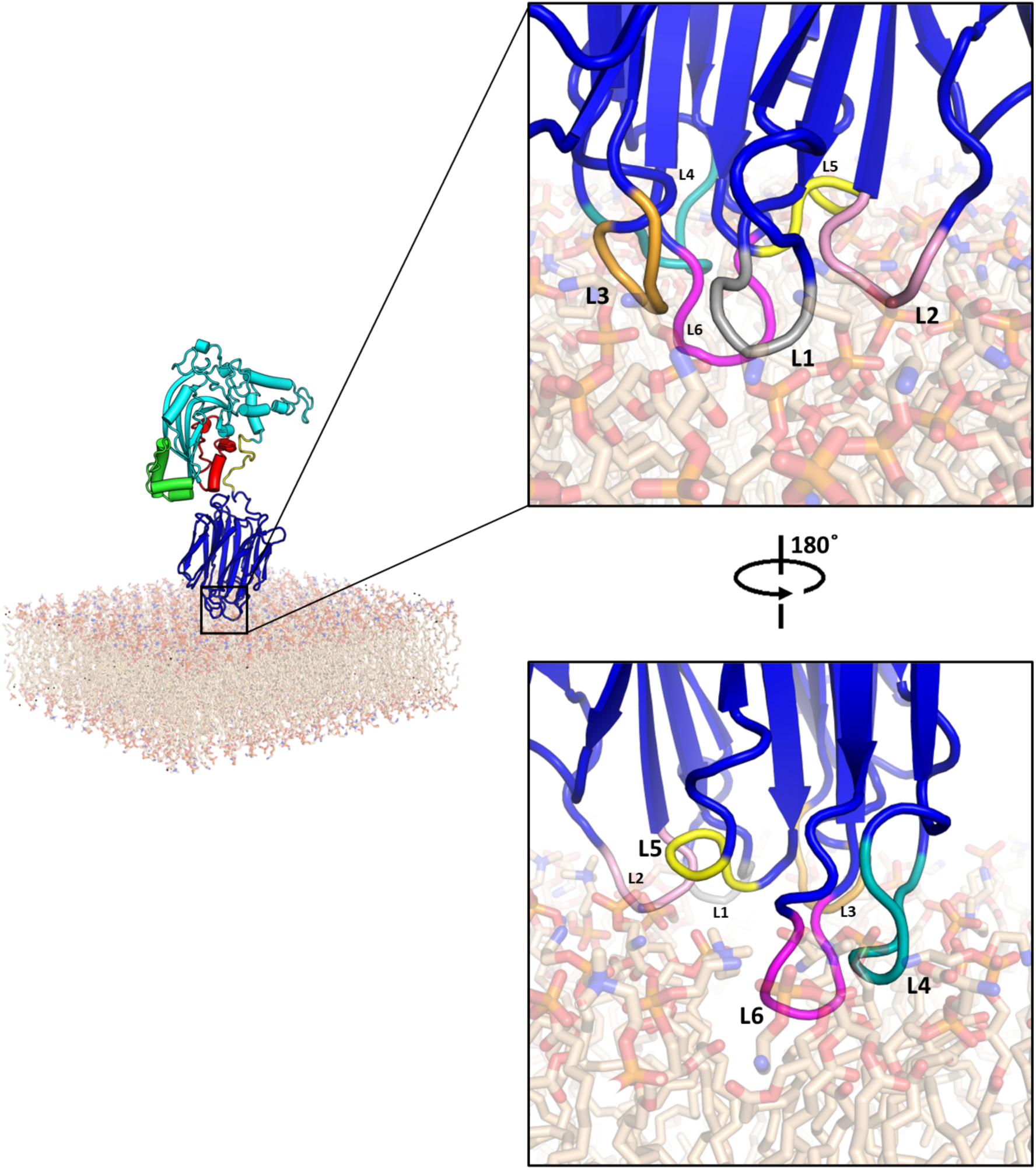
Interactions of PfPLP1 with the membrane mediated by the APCβ domain loops. Cartoon representation of the PfPLP1 monomer in soluble form interacting with the membrane. The zoomed-in insets present the 6 loops in different colors: L1-Gray (623-626), L2-Pink (651-654), L3-Orange (703-707), L4-Teal (764-770), L5-Yellow (797-800), and L6-Magenta (823-828). The lipid bilayer is represented as a transparent stick model.

The simulation trajectory of the soluble form of PfPLP1 on the lipid bilayer shows that the loops, L1 to L6, are involved in binding and interactions with the lipid bilayer; however, L2, L3, and L4 play a major role in the stable binding and interactions. The cationic residues, Lys-653 (L2), Arg-705 (L3), Arg-767 (L4), of different loop regions interact with the negatively charged head groups of the lipids with distances ranging from 2.8 to 3.3 Å and facilitate the binding of PfPLP1 to the lipid bilayer (Figure 4). The cationic residues at the particular position are conserved in all the malaria-causing *Plasmodium* species (Figure S3). Along with these residues, Lys-764 (L4) and Lys-770 (L4) are observed to form stable but transient hydrogen bonds (Figure S4). The hydrogen bond interactions of these residues with lipid molecules have been observed throughout the simulation trajectories, suggesting the importance of these APCβ-cationic residues in binding to the membrane. The binding of PfPLP1 in the current studies differs from that observed previously in TgPLP1 (Guerra *et al*., 2018), where the TgPLP1 APCβ domain is longer and thus forms an extended loop 6 (L6) compared to PfPLP1. The TgPLP1 has a tryptophan residue, which is responsible for its binding to the membrane; however, Trp is replaced by Phe at this position in PvPLP2 but is missing in PfPLP1 due to a shortened APCβ domain. Although the binding domains of MAC, perforin1, and perforin 2 are structurally different from the PfPLP1 APCβ domain but they show binding to the membrane through cationic-mediated interaction (Menny *et al*., 2018; Pang *et al*., 2019; Ni *et al*., 2020).

**Figure 4:**
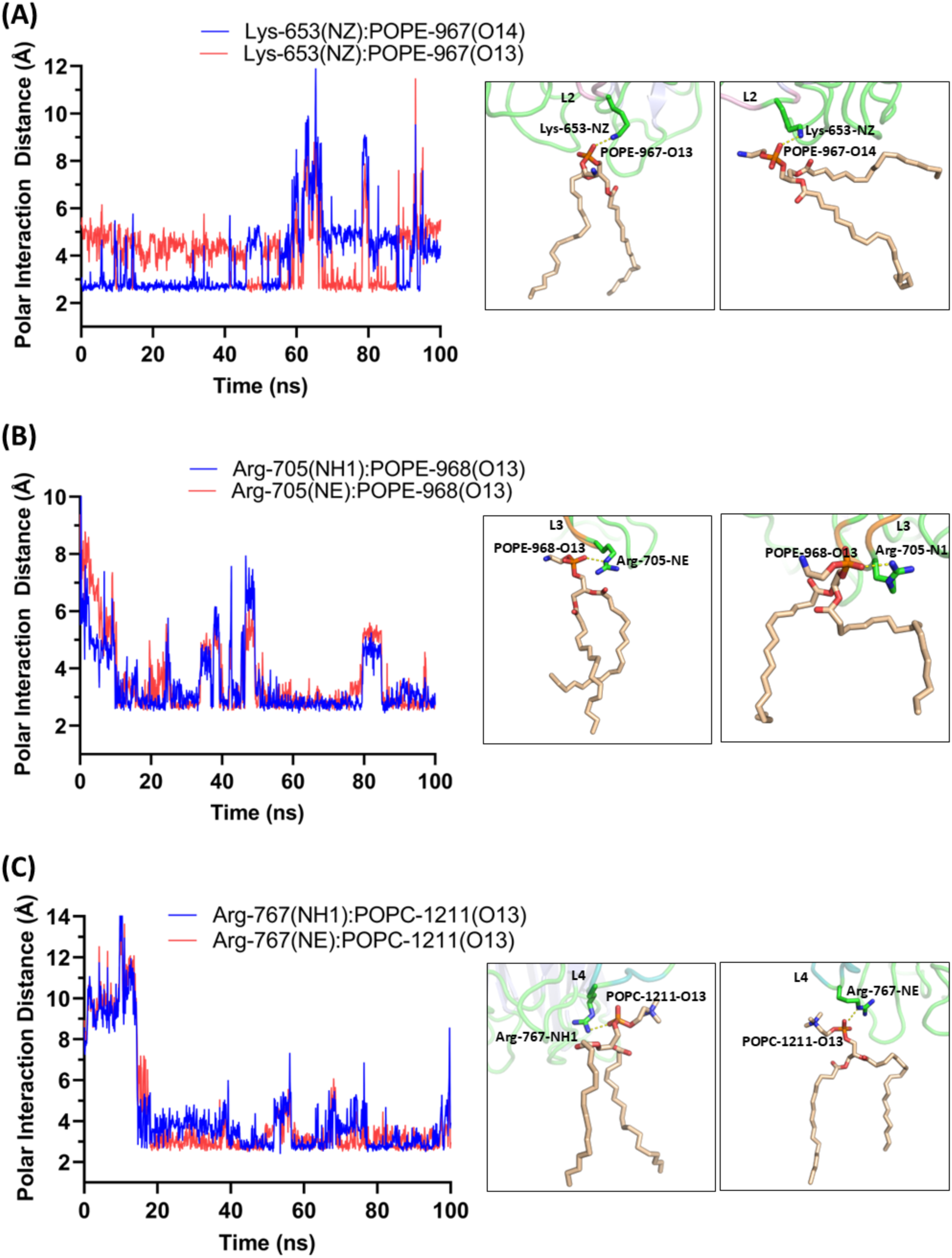
The polar interaction distances from the lipid molecules for PfPLP1 monomer on the lipid bilayer. (A) The Lys-653 of L2, (B) Arg-705 of L3, and (C) Arg-767 of L4 form stable interactions with the lipid head groups. The inset shows the interacting residues (green carbon) and lipid atoms (wheat carbon).

### All-atom MD simulations of tetrameric membrane-inserted PfPLP1

The interaction of the lipid bilayer and the membrane-inserted tetrameric form of PfPLP1 was analysed through all-atom MD simulations. The RMSD values for the tetramer system range between 0.45-0.8 units. RMSD vs Rg plot shows that the PfPLP1 tetramer maintained a stable conformation through the simulation trajectory (Figure 5A). The tetramer forms an arc with two distinctive faces, an inside face towards the lumen and an outside face opposite (Figure 5B). The inside face formed by the tetramer, which interacts with the solvents, has a polar charge, and the outside face has a neutral charge, which facilitates the interaction with hydrophobic lipids (Figure 5C). The β-sheets facing the lipid tails are lined with hydrophobic residues (Figure 5C right), helping to anchor and stabilize the β-sheet region within the membrane environment, and the pore side is lined by charged residues (Figure 5C left), promoting water and ion conduction through the protein interior.

**Figure 5:**
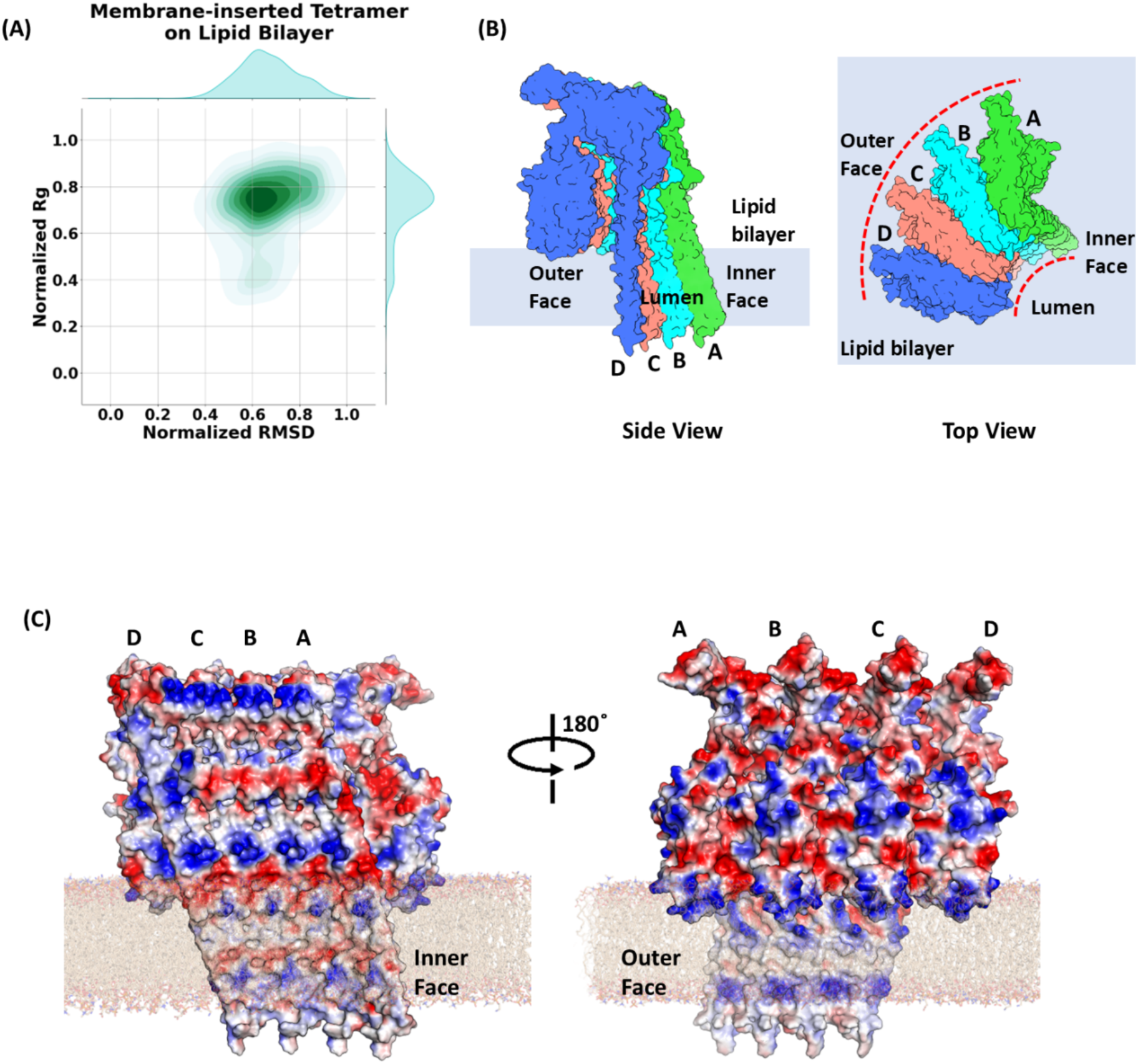
All-atom MD simulation of tetramer. (A) The Root Mean Square Deviation (RMSD) vs Radius of gyration (Rg) plot of the membrane-inserted tetrameric PfPLP1 during MD-simulation. (B) Schematic representation of the tetramer (each subunit colored with a different color: subunit A-Green; subunit B-Cyan; subunit C-Pink; subunit D-Blue) on the lipid bilayer (light blue rectangle). The lumen and the two faces, inner face and outer face, are represented in side view (left) and top view (right). The dotted line of the top view shows the formation of a small arc-like structure by tetrameric assembly. (C) The electrostatic surface charge distribution of the membrane-inserted PfPLP1 tetramer (blue: positive, red: negative, and grey: neutral) for the inner face (left) and outer face (right). The lipid molecules are represented as transparent sticks.

Initially, the polar residues present on the inside face of the β-sheets of the tetramer are in contact with the hydrophobic tails of lipid molecules (Figure 6A). Due to the charge distribution on the inside face, the hydrophobic lipid molecules are repelled away, which initiates the pore formation process within a few nanoseconds. Lipid molecules begin to reorganize themselves to shield the exposed hydrophobic region, and the reorganization begins as soon as 5 ns of the simulation (Figure 6B). However, the reorganizing is observed to occur throughout the simulation trajectory (Figure 6C), and by the end, a toroidal edge of the lipid bilayer is formed (Figure 6D), completely shielding the lipid tails and lipid head groups interacting with the polar residues and water molecules. During this reorganization, water molecules begin to insert between the β-sheets and the lipid molecules wetting the surface. Water penetration is observed from early simulation and observed throughout the trajectory, further stabilizing the polar residues on the inner face of the β-sheets (Figure 7). The water molecules also push the lipids away, further opening the pore. Similar pore opening and the toroidal edge formation of the lipids were also observed during MD-simulation studies on α-PFT, cytolysin-A from *E. coli* (Desikan *et al*., 2017a) and β-PFT, pneumolysin from *S. pneumoniae* (Vögele *et al*., 2019).

**Figure 6.**
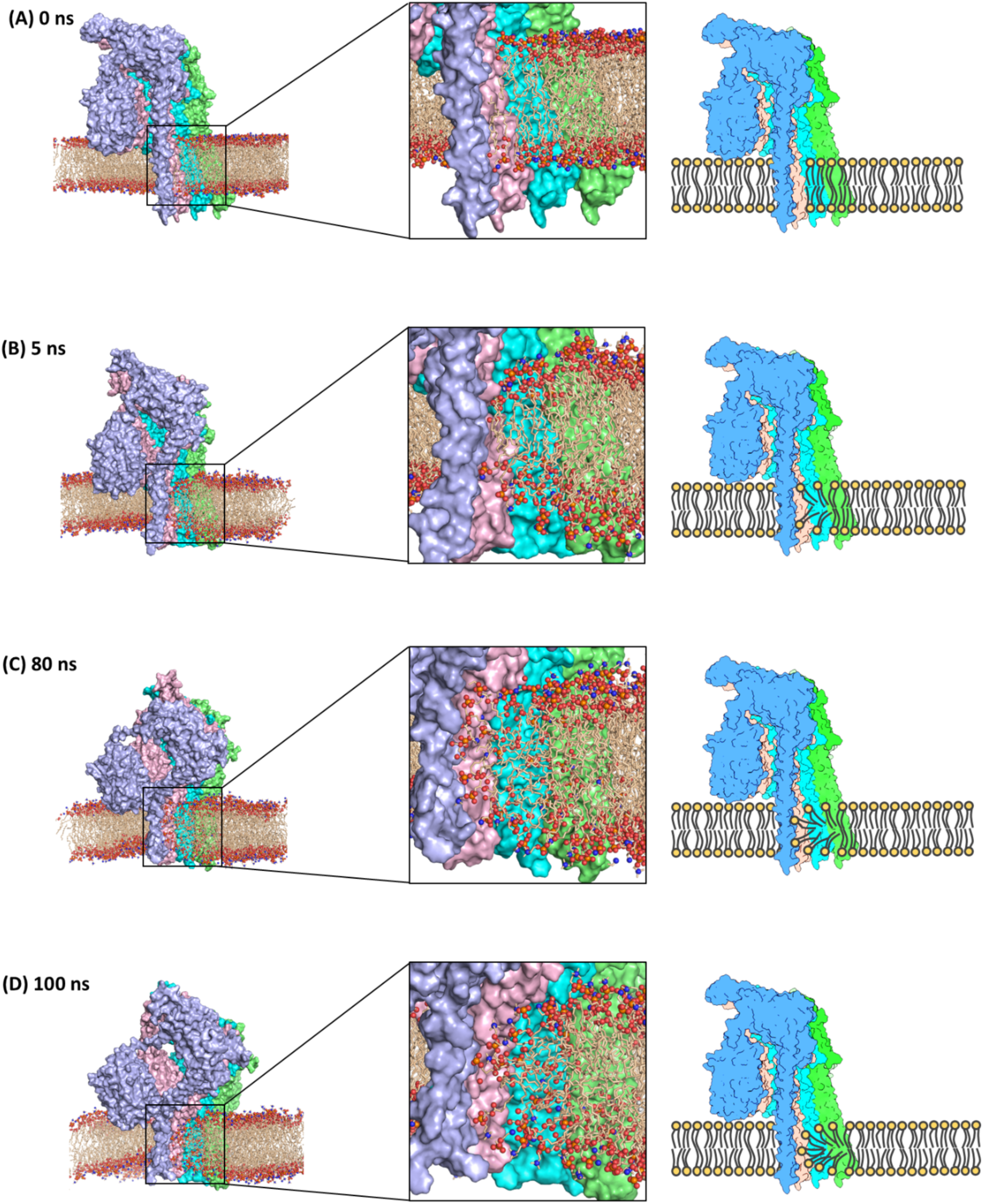
All-atom simulation of membrane-inserted PfPLP1 tetramer showing the toroidal edge formation of the lipids. The subunits in the system are shown in surface representation (subunit A in green, subunit B in cyan, subunit C in pink, and subunit D in light blue). The lipid molecules (represented in wheat colored sticks with oxygen and nitrogen atoms as red and blue balls, respectively) for 0 ns, 5 ns, 80 ns, and 100 ns frames. For better visual representation, water molecules and ions are removed from the frames. The lipids are exposed to the polar inner face (A). The reorganization of lipid molecules begins with the simulation (B). As the simulation progresses, the lipids form a toroidal edge shielding the lipid hydrophobic tails from the solvent (C and D). Schematics of this toroidal edge formation by PfPLP1 are also presented for better understanding on right side.

**Figure 7.**
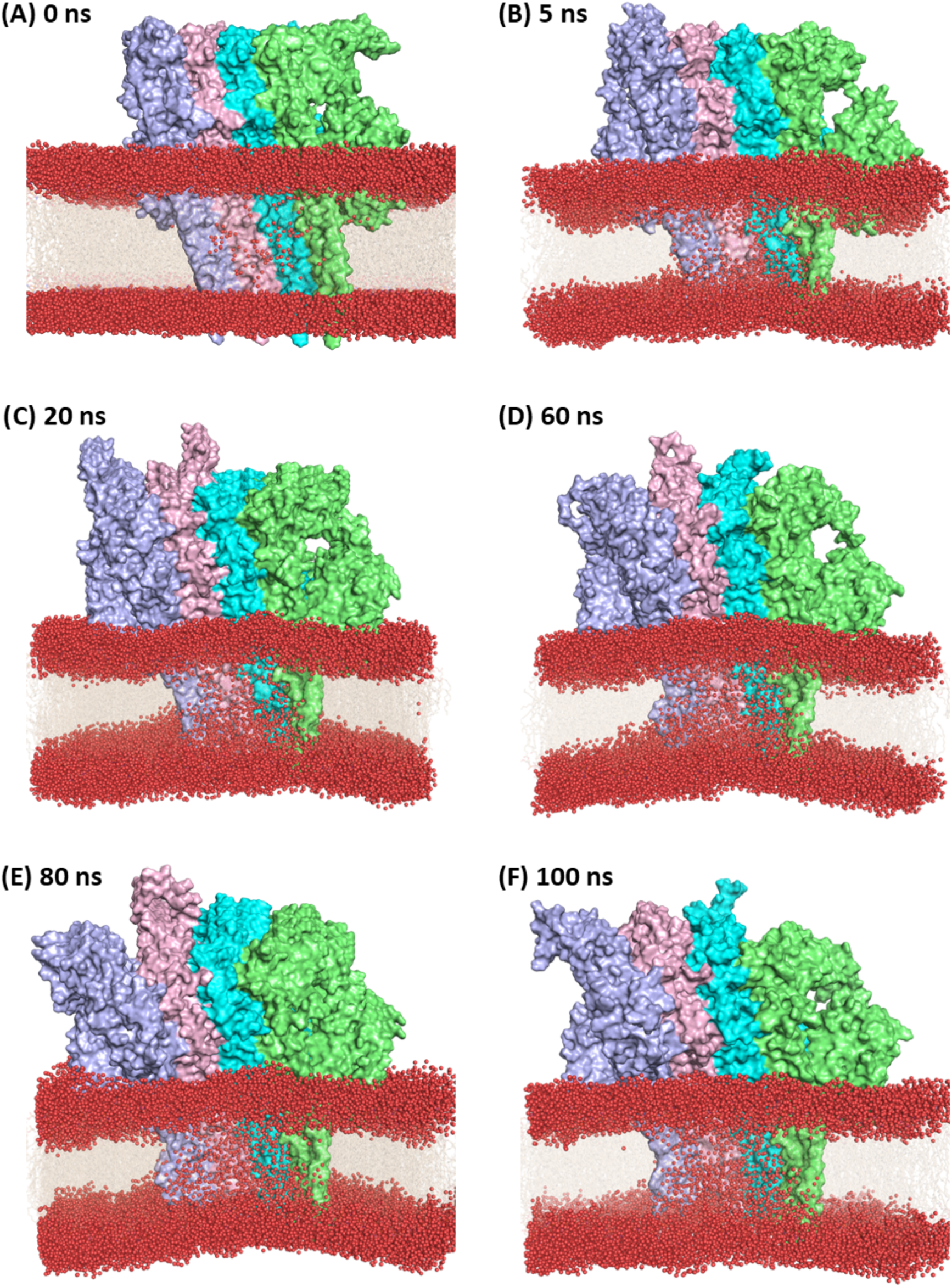
The water insertion and pore opening during 100 ns all-atom MD simulations for the tetramer of membrane-inserted PfPLP1. The subunits in the system are shown in surface representation (Subunit A in green, subunit B in Cyan, subunit C in pink, and subunit D in blue). The water molecules (red balls) present 10nm above and below the lipid bilayer (represented by wheat-colored transparent sticks) are shown in 0 ns, 5 ns, 20 ns, 60 ns, 80 ns, and 100 ns frames. For better visual representation, other water molecules and ions are removed from the frames. The water molecules start to move inside the protein arc and lipid molecules as soon as the simulation starts, increasing with the simulation time, in turn pushing the lipids away from the protein and opening the pore.

In membrane-inserted forms, the β-sheets are formed from structural rearrangement of TMHs and are very stable due to intra- and inter-chain hydrogen bond interactions. It is observed that one subunit of membrane-inserted PfPLP1 forms stable main-chain hydrogen bonds with the neighbouring subunit. The inter-chain hydrogen bond distances between the amino acids of neighbouring β-strands are mostly within the range of 2.92 to 3.40 Å, as shown in Figure 8. The intra-chain hydrogen bonds are between 2.8 and 3.5 Å for most of the simulation trajectory (Figure S5, S6, S7). Again, these main chain hydrogen bond interactions are important for the stabilization of the pore and pore-like structures. Recent cryo-EM structures of pores formed by MACPF domain proteins and CDCs have been observed to have such interactions between the neighbouring β-strands of subunits (Ivanova *et al*., 2022; Yu *et al*., 2022; Marini *et al*., 2023; Johnstone *et al*., 2025;).

**Figure 8.**
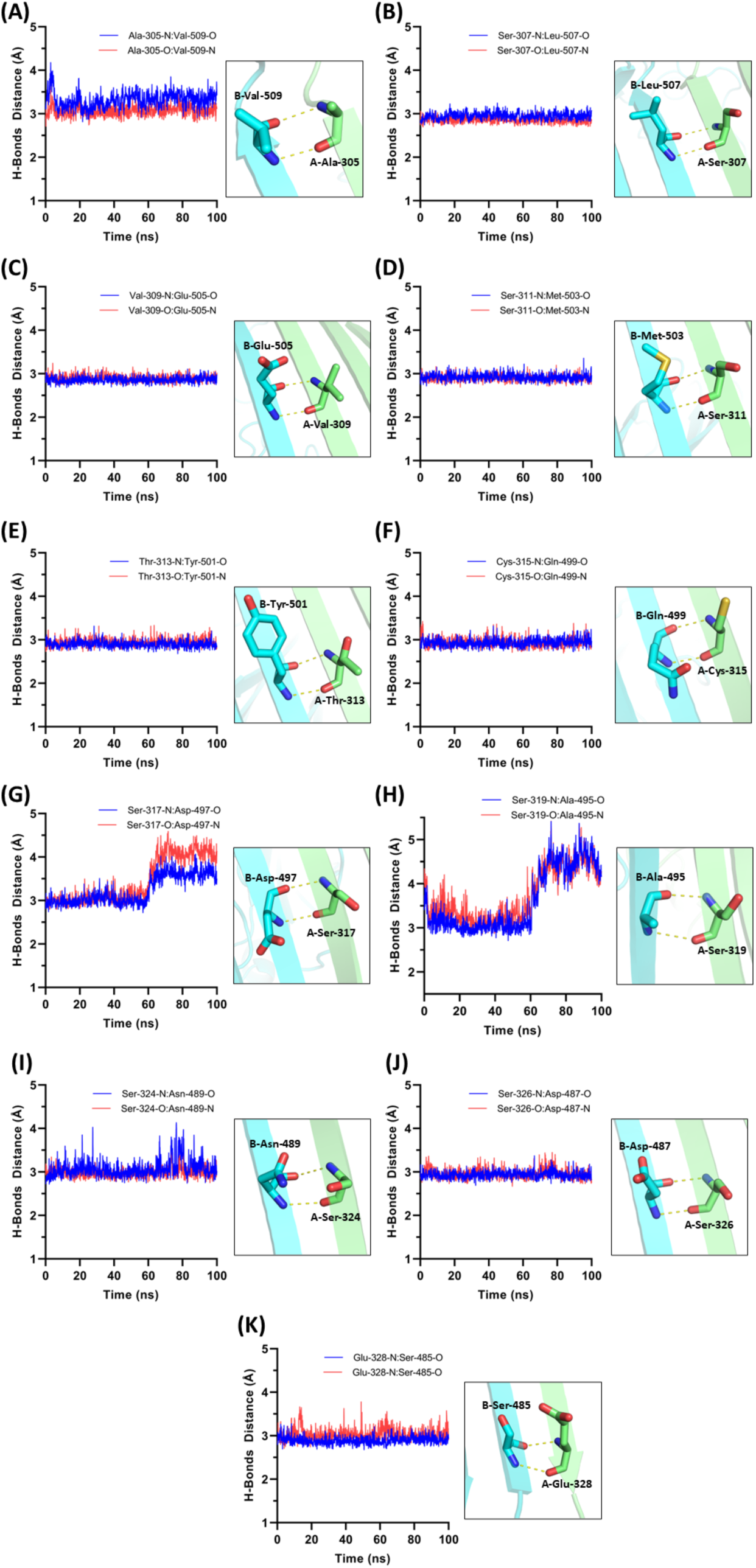
Inter-subunit hydrogen bond distances between different residues of neighbouring β-strands of two different subunits of the membrane-inserted tetramer of PfPLP1. The amino acid residues present in the adjacent β-strands of neighbouring subunits form stable main-chain hydrogen bonds. Inset: part of the Β-strands is shown as cartoon and the interacting residues are shown as sticks. The distances are shown as dotted lines.

### Structural analysis of PfPLP1 oligomers during coarse-grained (CG) MD simulations

We have performed CG MD simulation of the hexamer, octamer, decamer, dodecamer, and tetradecamer of membrane-inserted PfPLP1 for 1 µs to decipher the pore-forming mechanism and the fate of the lipids present within the pore. The analysis of the MD data revealed that oligomeric complexes are very stable throughout the simulation trajectory. The normalized RMSD versus Rg plots of the oligomeric complexes are shown in Figure 9, indicating their stability. Although the RMSD and Rg have higher fluctuation, they tend to stabilize by the end of the simulation. The higher fluctuation in RMSD is observed due to the tendency of the larger oligomers to form a pore-like structure, which is shown in Figure 10. The RMSD for hexamer (Figure 9A) and octamer (Figure 9B), where pore-like structure formation was not observed, ranges between 0.8-0.95 units, as the decamer (Figure 9C), dodecamer (Figure 9D), and tetradecamer (Figure 9E), which lead to the formation of pore-like structure, show higher fluctuation in RMSD, ranging between 0.6-0.98 units. All the time-dependent simulation curves are shown in the supplementary figure (Figure S8). The tendency of oligomeric arc to form a pore-like structure is observed in the α-PFT, Cly-A, from hexamer to larger oligomers like decamer. This tendency was believed to be due to the inherent property of PFTs to form pores (Desikan *et al*., 2017a; Desikan *et al*., 2017b). However, in our study, pore-like structure formation was not observed in smaller oligomers (hexamer, octamer) (Figure S9) (Supplementary Movies 1, 2), but with larger oligomers (decamer and above) (Supplementary Movies 3, 4, 5), which suggests that the formation of pore-like structures could be an inherent property of PFTs once a critical number of subunits are present.

**Figure 9:**
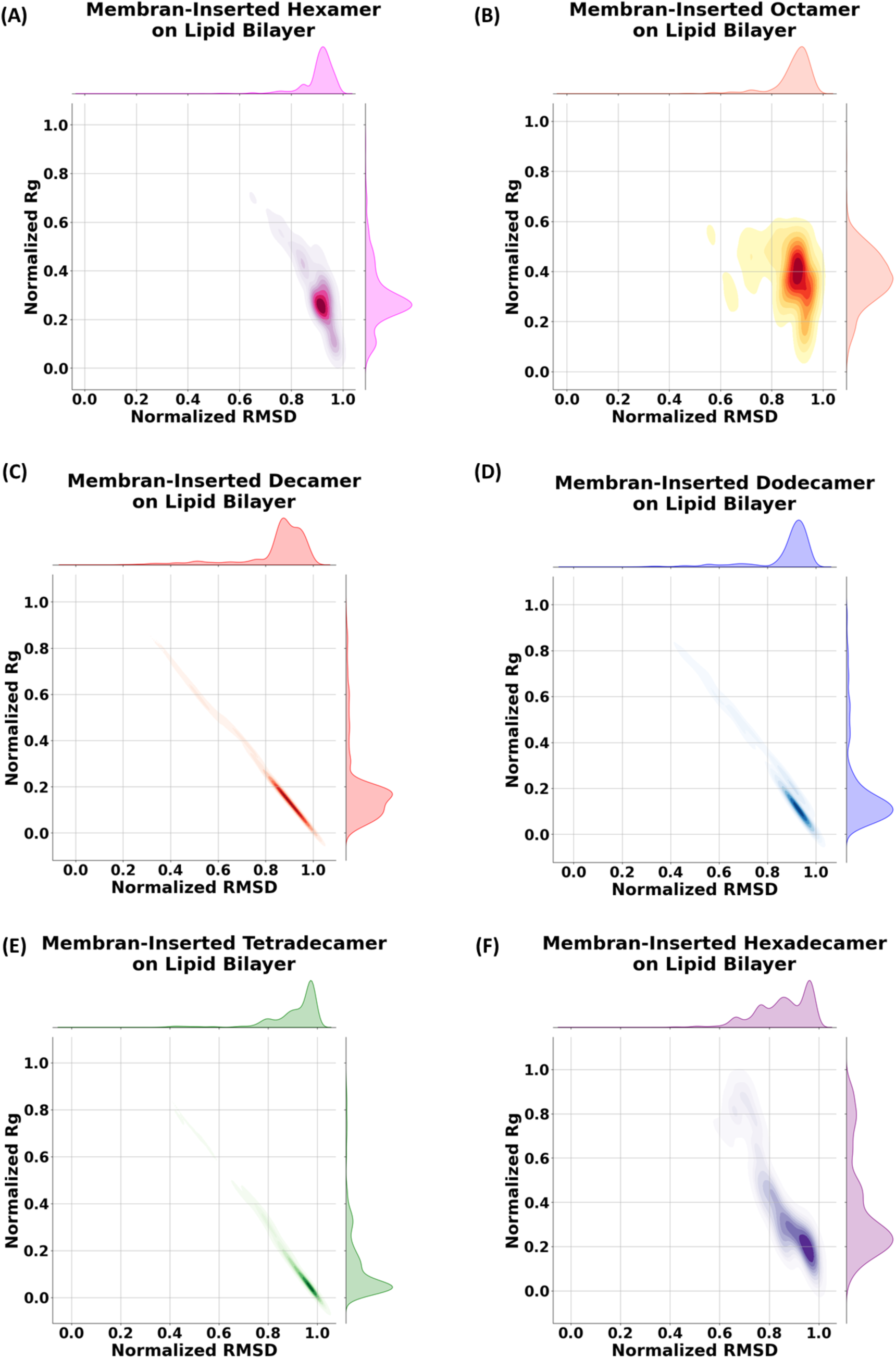
The Root Mean Square Deviation (RMSD) vs Radius of gyration (Rg) of the membrane-inserted oligomeric PfPLP during MD-simulations. (A) Hexamer (B) Octamer (C) Decamer (D) Dodecamer (E) Tetradecamer (F) Hexadecamer.

**Figure 10:**
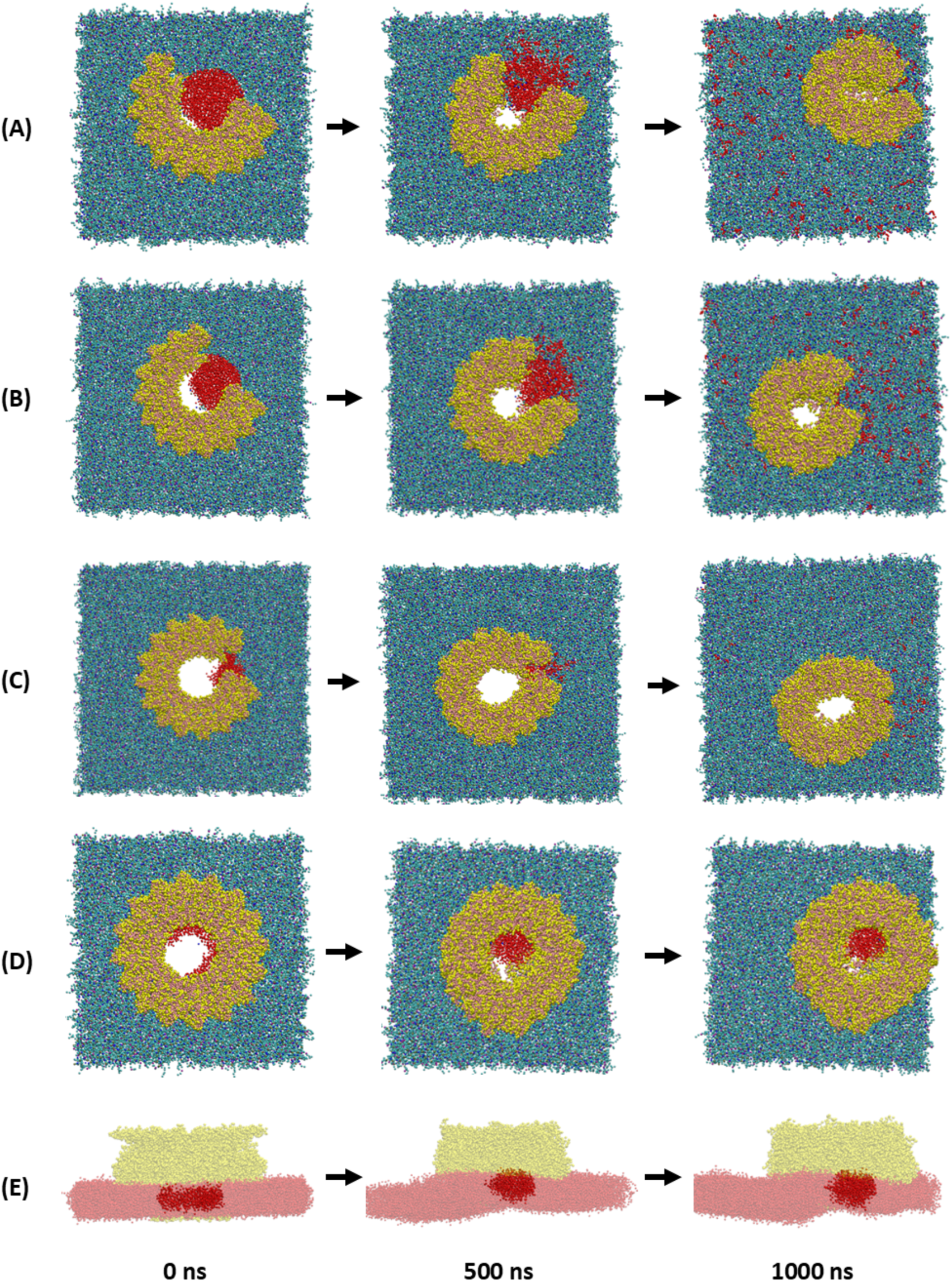
The fate of the lipids present inside the lumen of the oligomers during the simulations. Top-views of the pores formed by (A) Decamer, (B) Dodecamer, (C) Tetradecamer, (D) Hexadecamer. (E) Side view of the pore formed by Hexadecamer. The lipids present inside the arc of decamer, dodecamer, and tetradecamer are highlighted in red, which are excluded laterally as the simulation progresses. The lipids present inside the hexadecamer lumen form a micelle-like structure, which remains at the center of the lumen and moves slightly upwards. The lipids are highlighted in red and the protein in yellow. For the hexdecamer side-view, the lipids present outside the pore, also represented in red, and the proteins are made transparent for better visualization of the lipids present inside the lumen. The lipids and the proteins are represented as van der Waals radii. For better visual representation, the solvent and ions are not shown here.

The simulation of the complete ring of PfPLP1, which is made of sixteen subunits, was simulated for ∼3 µs. This complete ring is a representation of the pre-pore to pore transition pathway. The hexadecameric ring, although it remains intact, has also shown movement on the lipid bilayer, leading to higher fluctuation in the RMSD and Rg values (Figure 9F); however, it is comparably less than that of decamer, dodecamer, and tetratecamer (Figure S8). The diameter of the hexadecameric pore varies throughout the simulation, and the pore becomes smaller compared to the initial one, which is reported for MD-simulation studies of Cly-A and α-hemolysin (Desikan *et al*., 2017b).

The Atomic force microscopy (AFM) studies have indicated that the β-PFTs pore formation progresses through the pre-pore complex formation, followed by the transition of TMHs into β-sheets (Reboul *et al*., 2016). Formation of incomplete arcs were also observed by AFM in β-PFTs (Leung *et al*., 2014, 2017). During the formation of the membrane-inserted arc and the pore, the lipids present within the lumen are removed. The presence of the pore-like structure in incomplete arcs suggests a toroidal edge of lipids, shielding the hydrophobic tail from the water channel. Here, we have also analyzed the pore extension and lipid movement in the CG simulations of oligomeric PfPLP1 inserted in the membrane. The oligomers represent the arcs and incomplete rings, similar to those observed in AFM studies. The lipids present inside the arc lumen move laterally out, and the oligomer tries to form a complete pore during the simulation. The curvature of the oligomer changes, and the lipids get pushed outside (Figure 10A, 10B, 10C). During the pore formation, the lipids present at the inside face of the pore move away from the polar inside face (Supplementary Movies 3, 4, 5), similar to the one observed in all atom MD simulation of the terameric membrane-inserted PfPLP1. This tendency to form pore-like structures and the lateral movement of lipids have not been seen in smaller oligomers (Figure S9), suggesting that a critical number of oligomers is required for it to form a pore. Formation of the toroidal edge of lipids is also seen in CG simulations. This leads to the formation of a lipid-depleted zone, widening the pore. The lateral movement of lipids has also been observed in the MD simulations of incomplete arcs formed by membrane-inserted Cly-A and Ply (Desikan *et al*., 2017a; Desikan *et al*., 2017b; Vögele *et al*., 2019), indicating the lipid expulsion pathway in the pores of incomplete arcs.

Pore formation by β-PFTs proceeds via a prepore intermediate. During the transition from prepore to pore, lipids become trapped within the pore lumen. While the precise fate of these lipids remained unclear, it is suggested that lipids form micelle-like structures inside fully formed rings and may eventually exit the pore. To investigate this, we simulated the hexadecameric membrane-inserted PfPLP1 pore. Our observations revealed that lipids trapped within the pore lumen assemble into a stable micellar-like structure centrally located within the hexadecameric ring, with no vertical escape from the lumen observed in the simulation. Although no structural evidence for micellar removal of the lipids is available, MD simulation studies with Cly-A and Ply have shown the micellar removal by vertical movements (Vögele *et al*., 2019; Desikan *et al*., 2020). In PfPLP1, we observed that larger oligomers (decamer, dodecamer, and tetradecamer) show removal of the lipids laterally and a tendency to form a pore-like structure. However, during the simulation of the hexadecameric membrane-inserted pore, we observed a slight vertical movement of the micelle (Figure 10D, 10E) (Supplementary Movies 6, 7), but not the complete escape of the micelle from the lumen, so vertical movement may be possible, but our results dominantly support lateral movement of lipid molecules.

### Conclusions

The growing threat of drug resistance necessitates the exploration of new drug targets for combating malaria. The *Plasmodium falciparum* perforin-like proteins (PfPLPs) are crucial for the invasion and egress of the parasite, and they are considered potential drug targets. However, there is no structural data available for any of the PfPLPs, which has made it difficult for the structure-based drug development studies against PfPLPs. To the best of our knowledge, this is the first study focusing on the structural analysis of the PfPLPs using MD simulations. We successfully modeled both the soluble and membrane-inserted forms of PfPLP1, including its tetrameric membrane-inserted structure, to explore the mechanism of pore formation. Similar to other MACPF (membrane attack complex/perforin) family proteins, PfPLP1 anchors to the membrane via cationic residues, primarily lysine and arginine. In its membrane-inserted form, the β-strands exhibit alternating polar and non-polar residues where the hydrophobic face interacts with lipid tails, and the polar face lines the pore, facilitating solvent interaction. In all membrane-inserted forms, the β-strands engage in both inter- and intra-molecular hydrogen bonding, contributing to structural stabilization within the lipid bilayer. Smaller oligomeric assemblies of membrane-inserted PfPLP1 exhibited partial pore opening and lateral widening. Although complete pore formation was not observed in these assemblies, the presence of solvent flow through the arcs suggests that water molecules and small ions are capable of permeating the membrane. Both atomistic and coarse-grained simulations revealed that water molecules infiltrate the charged inside face of the β-sheets, displacing lipid tails and promoting pore expansion. These findings highlight the dynamic nature of PfPLP1-mediated membrane disruption and show the potential of identifying and targeting key structural features such as binding grooves or pockets for the development of novel inhibitors through structure-based drug design.

## SUPPORTING INFORMATION

Supporting information can be found as a separate file.

## ABBREVIATIONS

MD: Molecular dynamics;
ns: nanosecond;
nm: nanometre;
µs: microseconds;
PLP: Perforin-like protein;
PFTs: Pore forming toxins;
MACPF: Membrane attack complex/Perforin;
CDCs: Cholesterol dependant cytolysins;
APCβ: Apicomplexan perforin β-domain (APCβ);
CG: Coarse grained;
TMHs: Transmembrane β-hairpins;
Cryo-EM: Cryo-electron microscopy;
AFM: Atomic force microscopy;
POPC: 1-palmitoyl-2-oleoyl-sn-glycero-3-phosphocholine;
POPE: 1-palmitoyl-2-oleoyl-sn-glycero-3-phosphoethanolamine;
POPS: 1-palmitoyl-2-oleoyl-sn-glycero-3-phospho-L-serine;
Ply: Pneumolysin;
Cly-A: Cytolysin A;
RMSD: Root mean square deviation;
RMSF: Root mean square fluctuation;
Rg: Radius of gyration

## ACKNOWLEDGEMENTS

SP, SD, and PB thank to Indian Institute of Technology Bombay for providing research facilities in the Department of Biosciences and Bioengineering. Computational support from the Spacetime high-performance computational (HPC) facilities at IIT Bombay is acknowledged. We also acknowledge the inputs and help from Dr. Pooja Kesari.

## AUTHOR CONTRIBUTION

SP and SD designed, performed the experiments, and analyzed the data. SP and SD wrote the first draft of the manuscript, and PB reviewed and edited the manuscript. PB conceived the idea as well as was in charge of the overall direction, planning, and supervision of the project.

## DATA AVAILABILITY

Data are available from the authors upon request.

## CONFLICT OF INTEREST

The authors declare no competing interests.

## Notes

### Competing Interest Statement

The authors have declared no competing interest.

