## Supporting Information for "Deciphering Membrane Pore Formation Mechanisms of *Plasmodium falciparum* Perforin-Like Protein 1 (PfPLP1)"

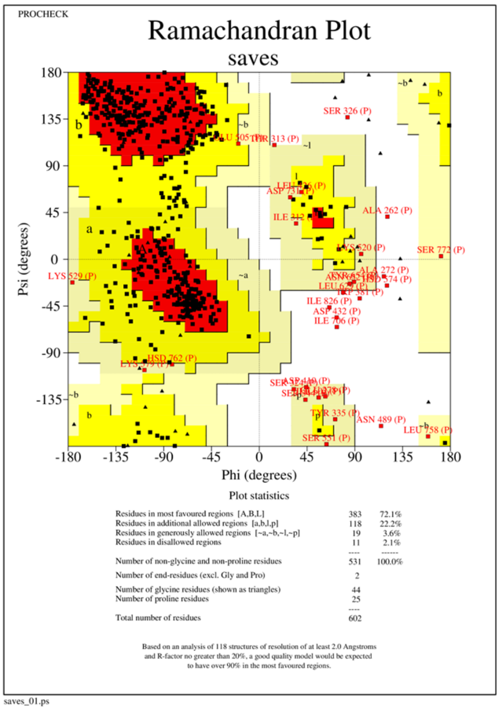


(B)

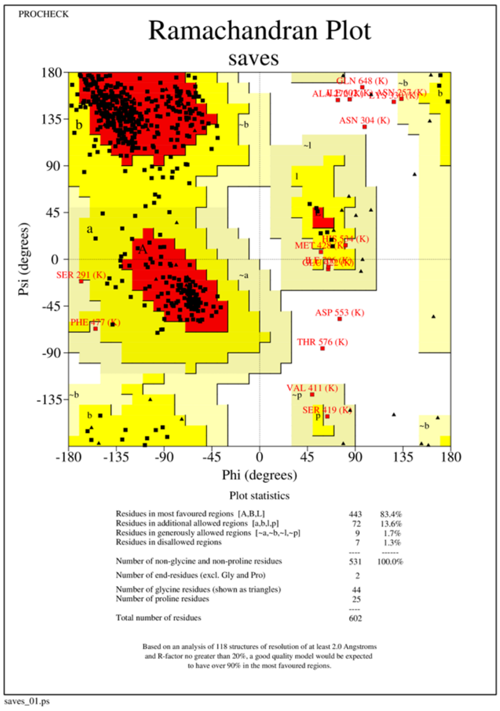


**Figure S1. The Ramachandran plots for the models generated for two different forms of PfPLP1.** (A) PfPLP1 in soluble form. (B) PfPLP1 in membrane-inserted form.

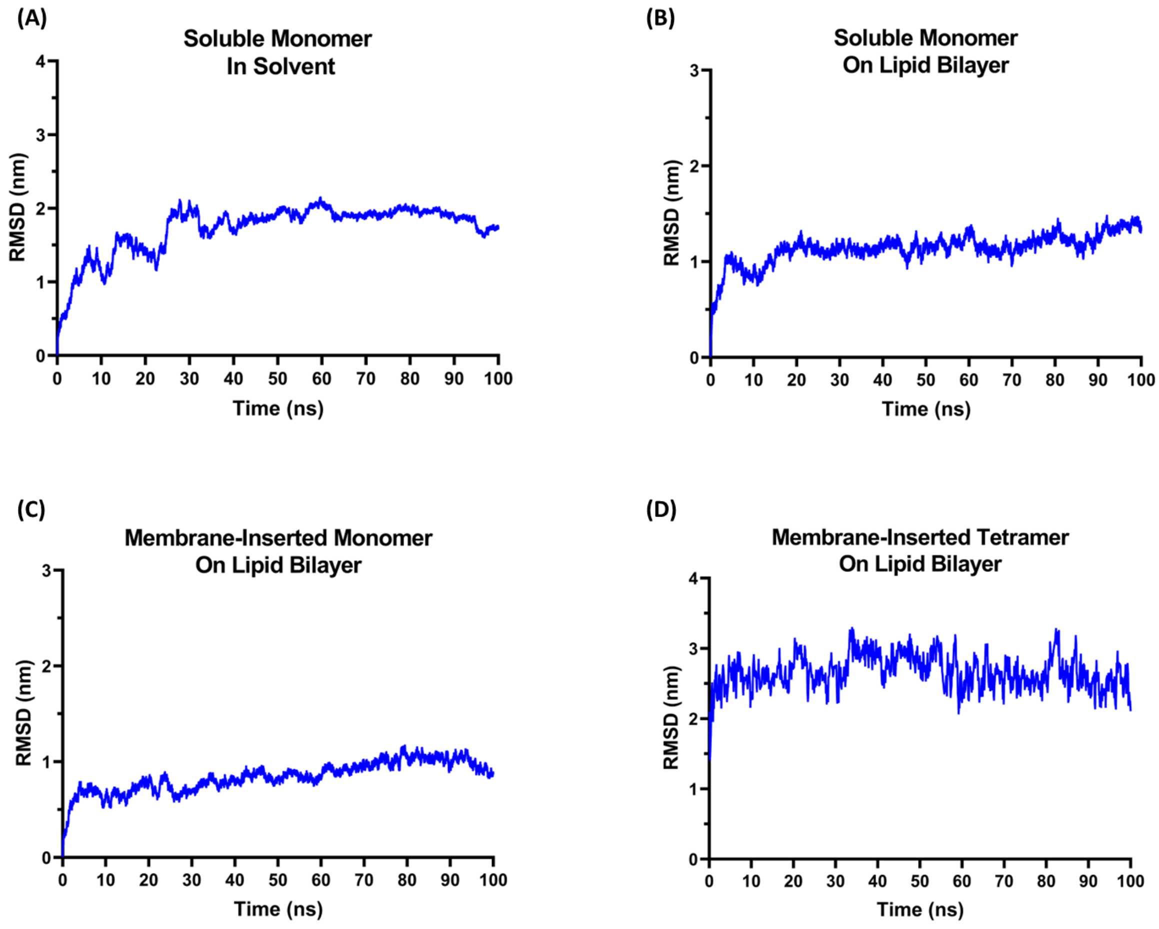


**Figure S2. RMSD plot of the all-atom MD simulation systems.** (A) Soluble PfPLP1 in solvent. (B) Soluble PfPLP1 on lipid bilayer. (C) Membrane-inserted PfPLP1 on lipid bilayer. (D) Tetramer of membrane-inserted PfPLP1 on lipid bilayer.

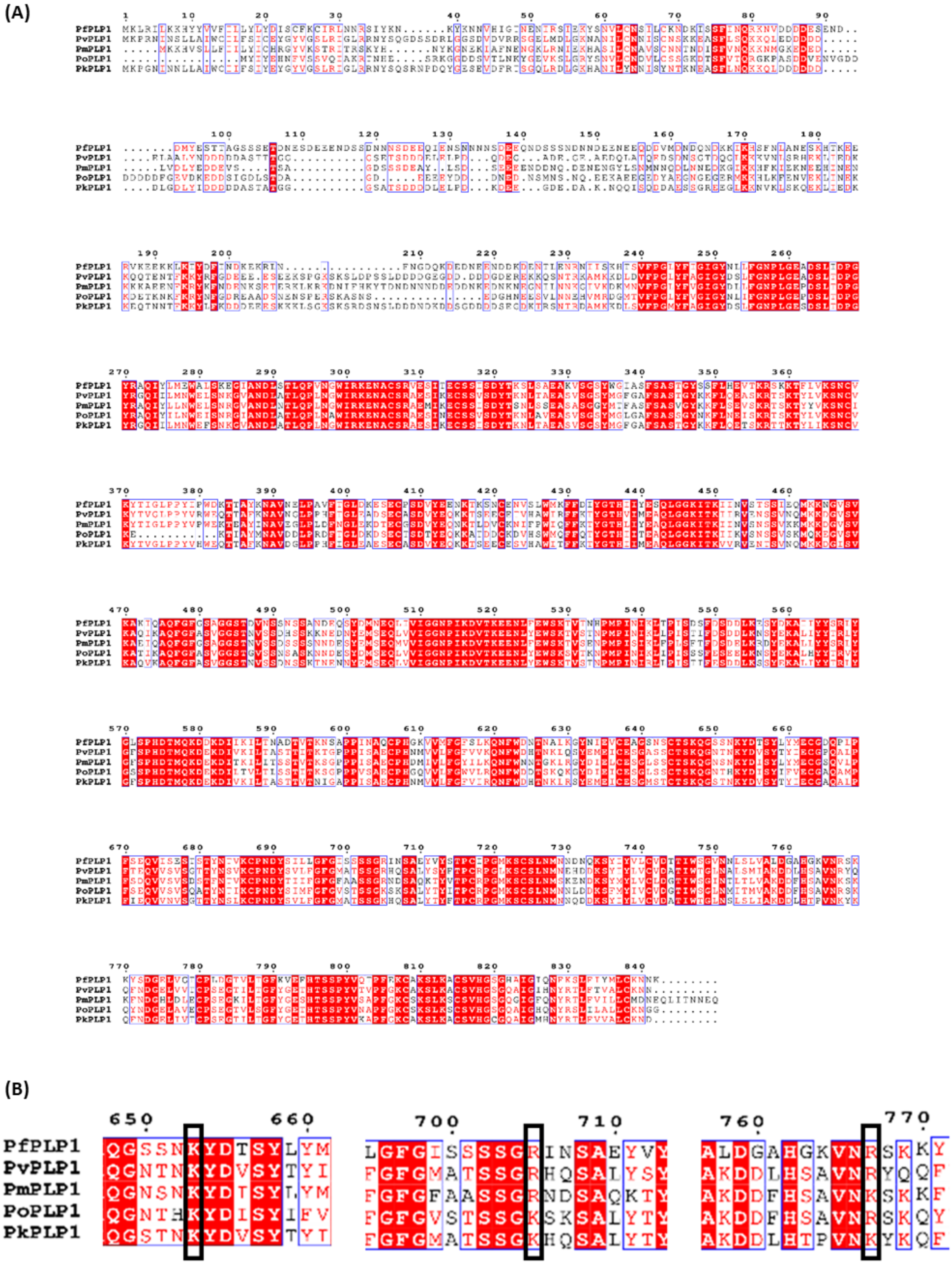

**Figure S3. Multiple sequence alignment of PLP1 from malaria-causing *Plasmodium* species.** (A) Complete sequence alignment of the full-length proteins. Residue numbers of PfPLP1 are also shown at the top of the alignments. (B) The conserved cationic residues among PLP1s are involved in the membrane binding. Pf: *Plasmodium falciparum*, Pv: *Plasmodium vivax*, Pm: *Plasmodium malariae*, Po: *Plasmodium ovale*, Pk: *Plasmodium knowlesi*.

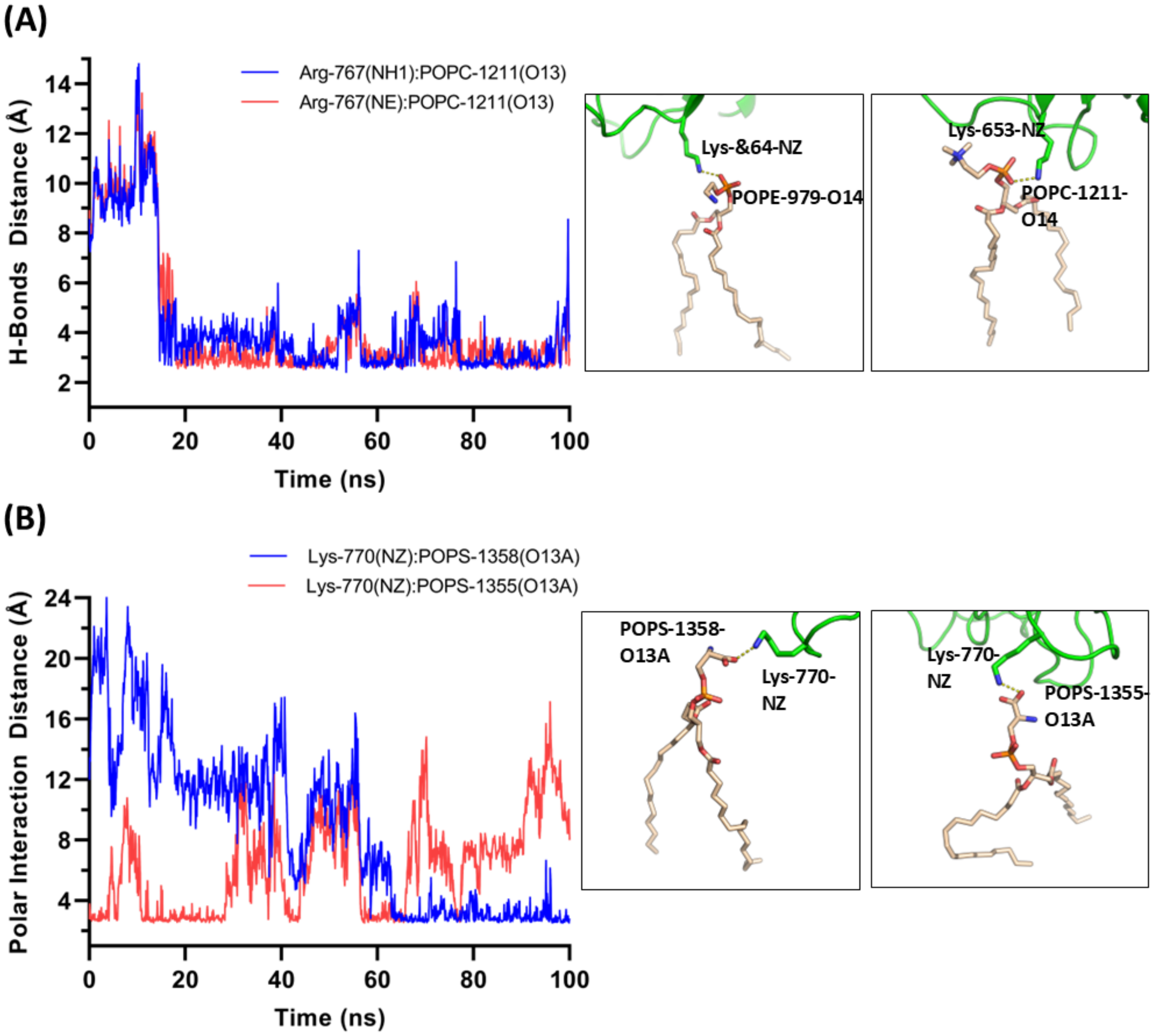


**Figure S4. transient polar interaction distances from the lipid molecules for the PfPLP1 monomer on the lipid bilayer.** (A) Lys746 residue of L4 and (B) Lys-770 residue of L4. The inset shows the interacting residues (green carbon) and lipid atoms (light brown carbon).

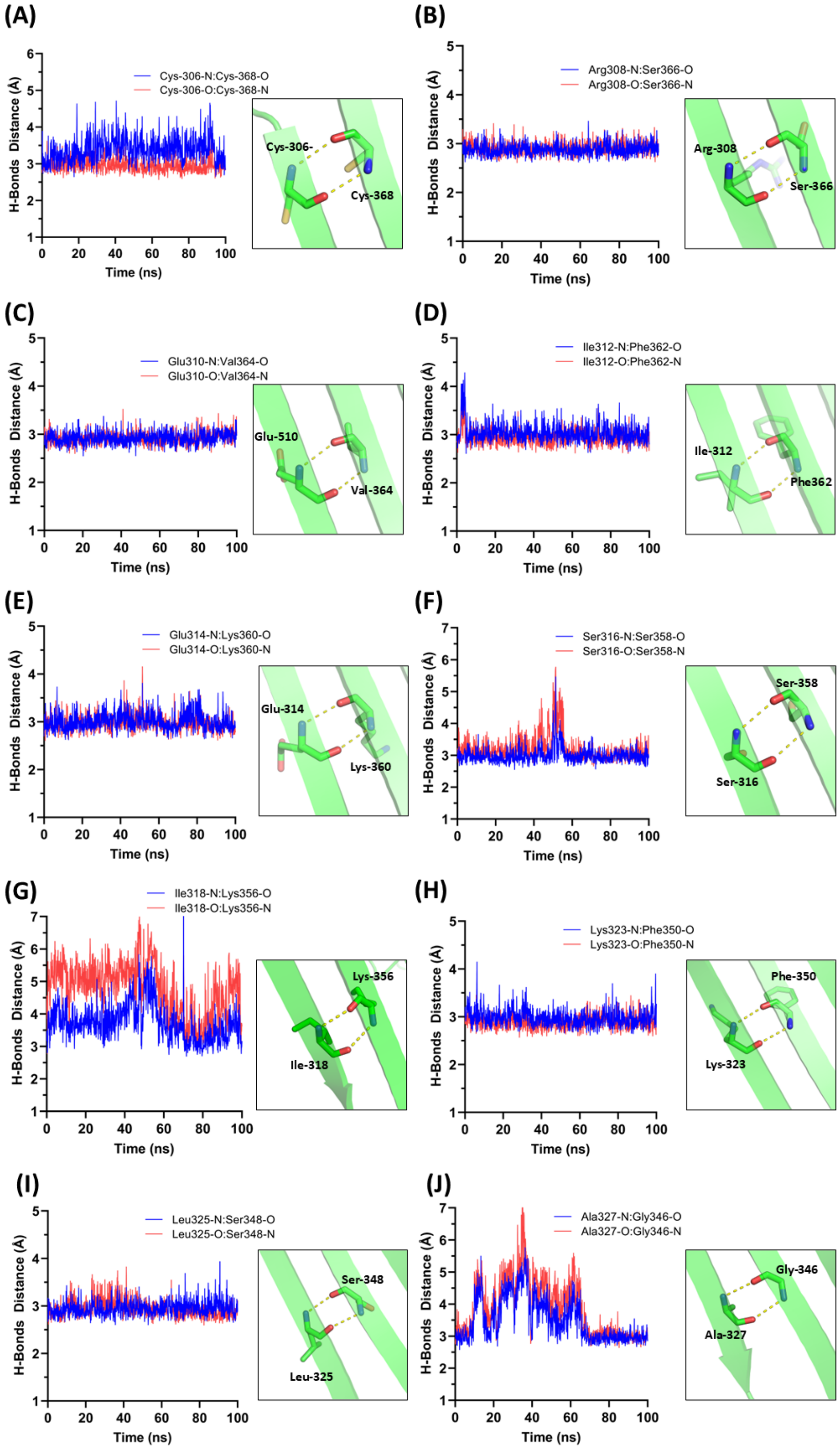


**Figure S5. Intra-subunit hydrogen bond distances between β-strand-1 and β-strand-2 of the membrane-inserted PfPLP1 in subunit A of the tetramer.** Inset: part of the β-strands is shown as a cartoon, and the interacting residues are shown as sticks. The distances are shown as dotted lines.

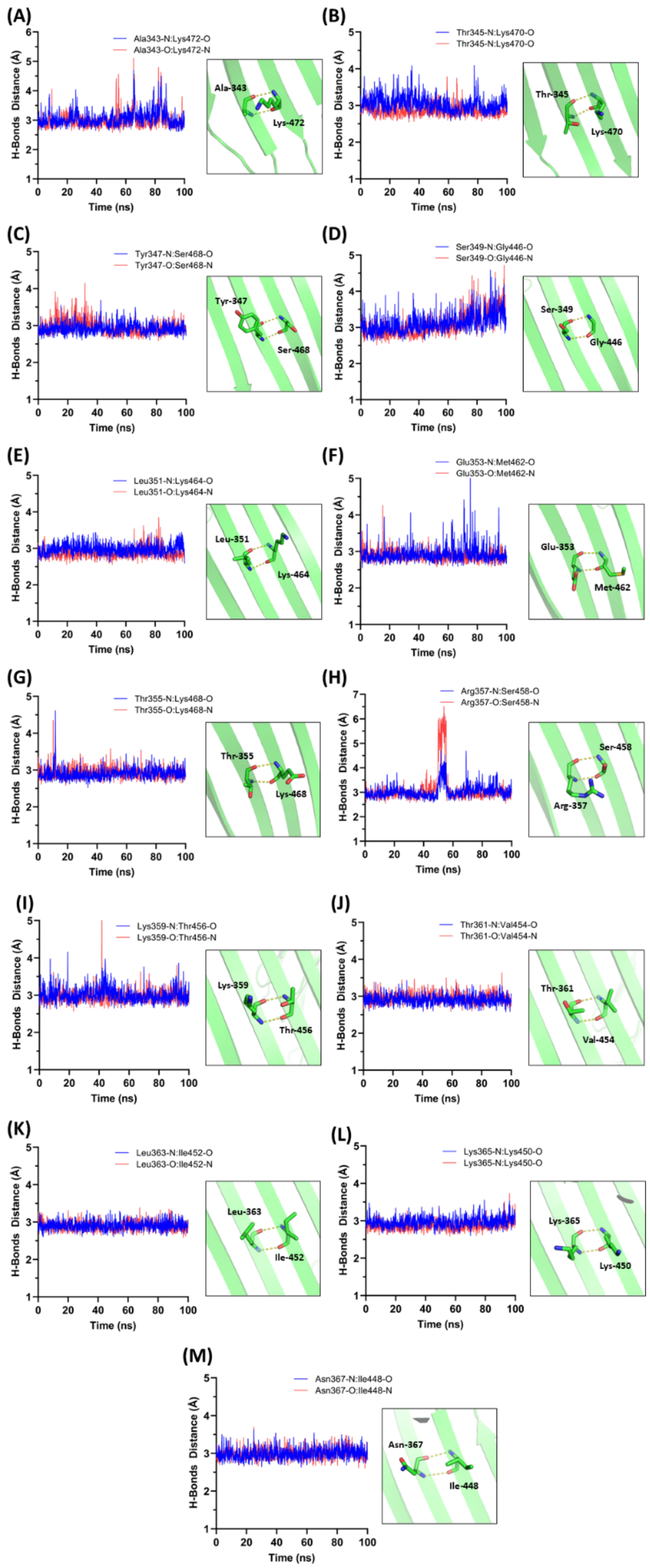


**Figure S6. Intra-subunit hydrogen bond distances between β-strand-2 and β-strand-3 of the membrane-inserted PfPLP1 in subunit A of the tetramer.** Inset: part of the β-strands is shown as a cartoon, and the interacting residues are shown as sticks. The distances are shown as dotted lines.

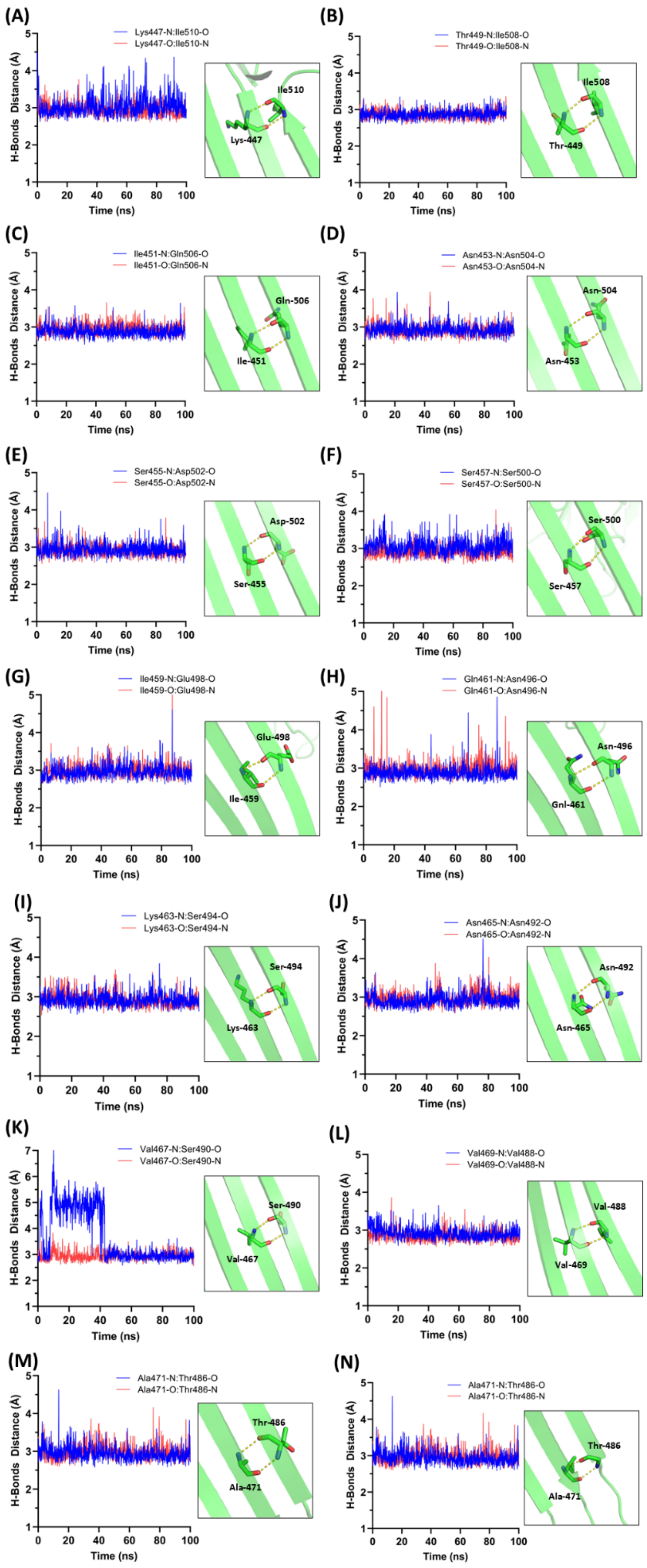


**Figure S7. Intra-subunit hydrogen bond distances between β-strand-3 and β-strand-4 of the membrane-inserted PfPLP1 in subunit A of the tetramer.** Inset: part of the β-strands is shown as cartoon and the interacting residues are shown as sticks. The distances are shown as dotted lines.

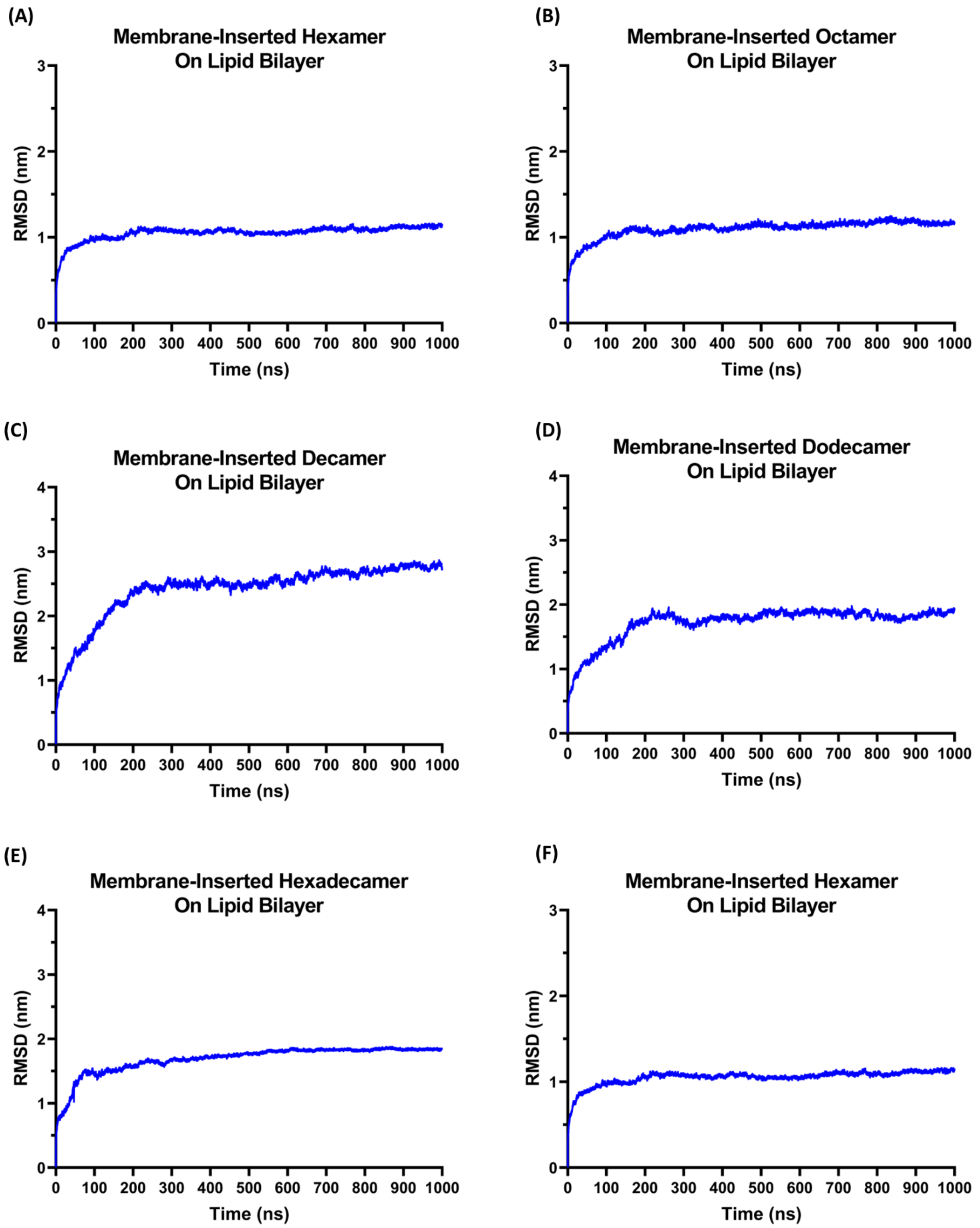


**Figure S8. RMSD plot of coarse-grained MD simulation systems.** (A) Hexamer (B) Octamer (C) Decamer (D) Dodecamer (E) Tetradecamer (F) Hexadecamer.

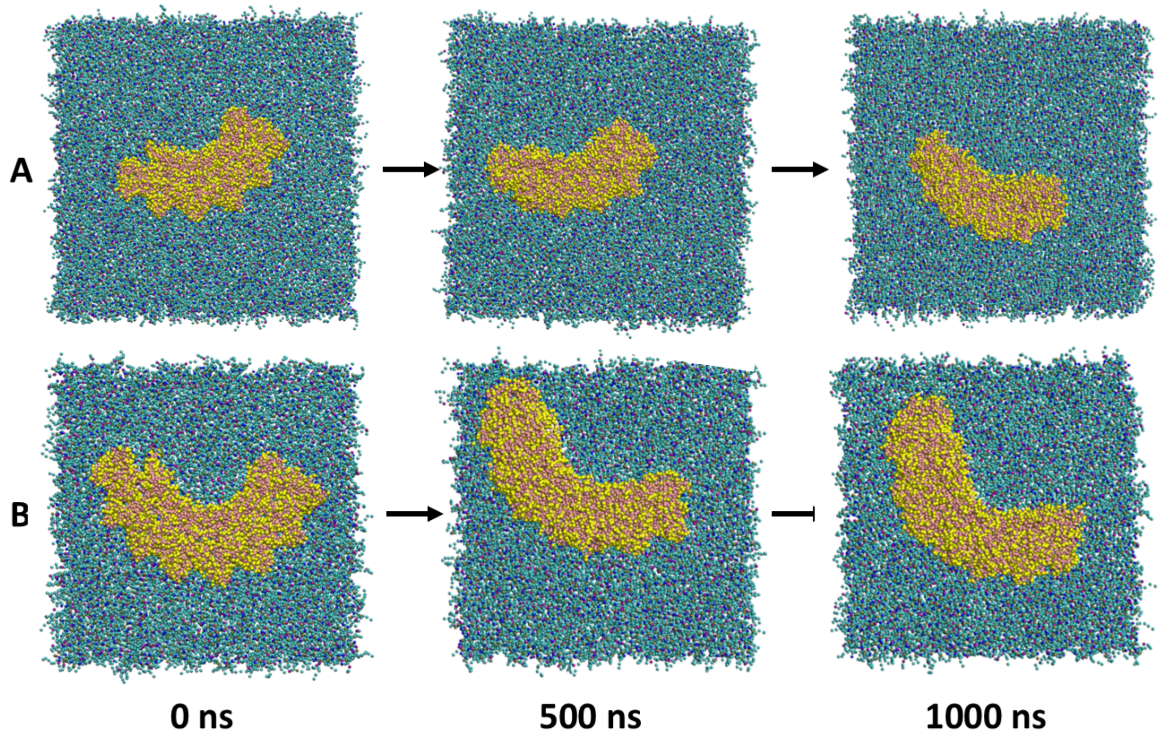


**Figure S9. The absence of a tendency to form pore-like structures in smaller oligomers.** (A) Hexamer (B) Octamer. The lipids and the proteins are represented as van der Waals radii. For better visual representation, the solvent and ions are not shown here. These oligomeric structures remain as arcs.

### Supplementary movie legends

**Movie 1: The CG-simulation trajectory of the membrane-inserted hexamer on the lipid bilayer.** The lipids and the proteins are represented as van der Waals radii. For better visual representation, the solvent and ions are not shown here.

**Movie 2: The CG-simulation trajectory of the membrane-inserted octamer on the lipid bilayer.** The lipids and the proteins are represented as van der Waals radii. For better visual representation, the solvent and ions are not shown here.

**Movie 3: The CG-simulation trajectory of the membrane-inserted decamer on the lipid bilayer.** The oligomer forms a pore-like structure, and lipids are excluded laterally from the lumen. The lipids present in the lumen are highlighted in red. The lipids and the proteins are represented as van der Waals radii. For better visual representation, the solvent and ions are not shown here.

**Movie 4: The CG-simulation trajectory of the membrane-inserted dodecamer on the lipid bilayer.** The oligomer forms a pore-like structure, and lipids are excluded laterally from the lumen. The lipids present in the lumen are highlighted in red. The lipids and the proteins are represented as van der Waals radii. For better visual representation, the solvent and ions are not shown here.

**Movie 5: The CG-simulation trajectory of the membrane-inserted tetradecamer on the lipid bilayer.** The oligomer forms a pore-like structure, and lipids are excluded laterally from the lumen. The lipids present in the lumen are highlighted in red. The lipids and the proteins are represented as van der Waals radii. For better visual representation, the solvent and ions are not shown here.

**Movie 6: The CG-simulation trajectory of the membrane-inserted hexadecamer on the lipid bilayer in top-view.** The oligomeric ring traps lipids inside the lumen. The lipids from a micelle-like structure stay inside the lumen. The lipids present in the lumen are highlighted in red. The lipids and the proteins are represented as van der Waals radii. For better visual representation, the solvent and ions are not shown here.

**Movie 7: The CG-simulation trajectory of the membrane-inserted hexadecamer on the lipid bilayer in side-view.** The micelle-like structure moves slightly upward in the lumen. The lipids are highlighted in red, and the lipids outside the lumen and the protein are made transparent for better visual representation of the lumen lipids. The lipids and the proteins are represented as van der Waals radii. For better visual representation, the solvent and ions are not shown here.
